# Maternal E-cigarette Exposure Augments Murine Abdominal Aortic Aneurysm

**DOI:** 10.64898/2026.10.19.760180

**Authors:** Joshua M. Spin, Joscha Mulorz, Wen-Jun Shen, Maryam Amirahmadi, Y. Hyun Rhee, Alicia Deng, Sohil Chavan, Jackie Butterfield, Pireyatharsheny Mulorz, Markus U. Wagenhauser, Colwyn A. Headley, Tao Qiu, Philip S. Tsao

## Abstract

**Background:** Smoking and family history are major risk factors for abdominal aortic aneurysm (AAA). Smoking and nicotine are modulators of DNA methylation, capable of causing heritable epigenetic alterations. Nicotine infusion and e-cigarette (e-cig) vaping augment murine AAA. We investigated the effects of murine maternal e-cig vaping on offspring experimental AAA, and impacts on transgenerational DNA methylation and accessibility.

**Methods:** We performed perinatal e-cig nicotine vaping in two strains of adult female mice (ApoE-/- and C57BL/6J). Mice were mated to control males, and treated through gestation. A parallel group were instead sacrificed after treatment, for DNA analysis. Offspring were aged to adulthood, and AAAs were then induced by elastase (C57BL/6J) or angiotensin-II infusion (ApoE-/-), and ultrasound-tracked for 4 weeks. In a second cohort, parental and offspring aortic tissue and blood DNA underwent RRBS and ATAC-seq analysis respectively. Offspring abdominal aortic anatomy was examined, and stiffness was assessed with ex vivo arterial myography.

**Results:** Maternal e-cig exposure increased offspring AAA growth in both models and genders. AAA incidence was increased in ApoE-/- males, and observed effects were overall more pronounced in males. Maternal vape exposure altered aortic DNA methylation and blood DNA accessibility in mothers and offspring, with differential sites in numerous genes identified, including AAA-related genes and pathways. Pathways were enriched for transcription factors, inflammation/immune-related genes, and angiogenesis genes, with similar changes identified in exposed mothers. Male offspring displayed more differential genes. Significant changes were located in genes known to harbor AAA risk loci in humans, and overlapped with published epigenetic alterations in human smokers. Differences also occurred in offspring aortic structure and stiffness.

**Conclusions:** Maternal e-cig nicotine exposure augments AAA growth in multiple models across generations. These effects are accompanied by broad epigenetic DNA reprogramming of key AAA modulator genes and pathways.

## Introduction

Abdominal aortic aneurysm (AAA) is defined as a pathologic dilatation of the aortic diameter > 30 mm, most often occurring in the infrarenal segment.^1^ AAAs cause significant morbidity and mortality, with the most feared clinical consequence being acute rupture.^2^ AAA pathology is complex, and involves local inflammation with leukocyte and macrophage recruitment, accompanied by elastin fragmentation and extracellular matrix degeneration, with phenotypic modulation and loss of medial vascular smooth muscle cells (SMC).^3^

Among the risk factors for AAA development are family history, and cigarette smoking, Further, smoking is the most important known modifiable risk factor for AAA; nearly all AAA patients (>90%) relate a smoking history.^4^ Ongoing smoking leads to more rapid aneurysm expansion,^5^ and cessation is the only treatment which clearly decreases the rate of AAA growth.^6^ While traditional cigarette use has been declining in Western countries, alternative nicotine delivery methods have increased in popularity. Introduced in 2003, electronic cigarette (e-cig) vaping has become prevalent in all ages, sexes and races, serving as a supposedly “healthier” substitute for smoking while delivering enough nicotine to satisfy addiction.^7^ Nicotine is a potent stimulant and parasympathetic-activating alkaloid. E-cigs and similar devices release “vapor” produced by heating a solution (known as “e-liquid” or “e-juice”), usually containing nicotine along with propylene glycol and glycerol, with or without flavoring chemicals.^8,9^

Several studies have associated e-cig use with elevated cardiovascular risk.^10–14^ A recent consensus paper highlighted the cardiovascular toxicity of nicotine, irrespective of delivery system, and described an urgent need for regulation.^15^ Previous studies show that both subcutaneous infusion of nicotine and e-cig vaping augment AAA expansion in murine models, altering cytokine production, triggering inflammation, and affecting vascular SMC phenotype.^16–19^ Nicotine infusion also induces aortic remodeling and regional vascular stiffness in humans and mice, which can predispose to AAA.^20–23^

However, both smoking and nicotine are also known to be potent epigenetic DNA modulators, inducing changes that may be durable for decades^24–28^. Accumulating evidence links maternal tobacco exposure to adverse vascular outcomes in offspring, potentially through durable alterations in epigenetic programming.^29–31^ Notably, 2.2%– 7.0% of pregnant women report e-cig use during pregnancy, numbers likely to be underestimates.^32^

It is therefore plausible that part of the heritable risk of AAA could be related to parental nicotine exposure. We explored the relationship between maternal e-cig vaping exposure and AAA-related risk and processes in their offspring in two mouse models.

## Methods

### Mice

All animal protocols were approved by the Administrative Panel on Laboratory Animal Care at Stanford University (https://researchcompliance.stanford.edu/panels/aplac) and the VA Palo Alto Health Care System Institutional Animal Care and Use Committee (protocol # SPI1933) and followed the National Institutes of Health and U.S. Department of Agriculture Guidelines for Care and Use of Animals in Research. All experiments were performed with C57BL/6J and ApoEtm1Unc (ApoE-/-) mice. Animals were purchased from The Jackson Laboratory (Bar Harbor, ME, USA).

### E-cigarette vapor exposure

At 8 weeks-of age, female (F0 generation) C57BL/6J and ApoE-/- mice were exposed to e-cig vapor containing 50% propylene glycol/50% vegetable glycerol (PG/VG), and 24 mg/mL nicotine using the SciReq InExpose system (SCIREQ, Montreal, Canada) using previously described protocols.^18^ The protocol included a puff duration of 9 s per minute, for a total of 60 min at the same time daily. The device used was a third-generation MOD e-cig (Shenzhen Joyetech Co., Shajing Town, China) with a temperature-controlled coil set to 230°C. Plasma cotinine levels were tested immediately after treatment in test mice via ELISA according to the manufacture’s protocol (Abcam, Cambridge, UK). Control animals were placed in SciReq inhalation chambers for the same daily duration but were exposed to room air only. Mice were either exposed to e-cig or room air for a total of 4 weeks and then harvested, or treated for 1 week, and then mated to same-strain control males, followed by treatment through gestation. Offspring (F1 generation) were aged to 10-12 weeks in room air, and then were either harvested for DNA analysis or ex vivo arterial myography, or underwent AAA induction.

### Porcine pancreatic elastase infusion model and angiotensin II (AngII) AAA models

The porcine pancreatic elastase (PPE) infusion model to induce mouse AAA was performed as previously described in both male and female C57BL/6J F1 offspring.^18,33^ The proximal and distal aorta were temporarily ligated or clamped, followed by an aortotomy above the iliac bifurcation. A catheter was used to infuse the aorta for 5 min at 120 mmHg with saline (‘sham’ surgery control), or saline containing type I porcine pancreatic elastase (2.5 U/mL; Sigma Aldrich), and the aortotomy was repaired. No deaths occurred during AAA development.

For the AngII model, ApoE-/- F1 offspring mice of both sexes were implanted subcutaneously with osmotic mini pumps (Alzet, Model 2004) filled with either sterile saline (control), or saline containing angiotensin-II (1µg/kg/min, Sigma Aldrich) as previously described.^18,34^ Deaths from spontaneous aortic rupture during the time-course were counted as positive for AAA incidence, although such events were not significantly different between controls and maternally vaped offspring offspring (FRA 8%, FVF1 11.11%, MRA 18.18%, MVF1 10%).

Prior to surgery mice received extended-release buprenorphine (Fidelis Animal Health Inc, NJ, USA) at 3.25 mg/kg subcutaneously. During surgery anesthesia was maintained with isoflurane 1.5-2% (Vet One, Meridian, ID, USA). Post-surgery, all animals received 2 drops of 0.25% bupivacaine locally before wound closure. Post-surgical daily monitoring was performed in accordance with the approved procedural protocols. Sacrifice was performed under surgery anesthesia by heart puncture. After flushing the aorta via the left ventricle with ice cold phosphate buffered saline (PBS; pH 7.4) it was dissected from fat and connective tissue from the left renal artery to the bifurcation using a microscope for the PPE model mice (Leica, Wetzlar, Germany). Blood and aorta were snap-frozen individually in liquid nitrogen and stored at −80°C before further processing for sequencing.

### AAA development ultrasound monitoring

AAA growth in both models was monitored as previously described via B-mode ultrasound (Vevo 2100® High-Resolution In Vivo Micro-Imaging System (VisualSonics, Toronto, Canada) at baseline and days 3, 7, 14, 21, and 28 post-induction.^18^ Mice were studied under isoflurane anesthesia and aortic diameters were assessed at maximum diameter in the systolic cardiac phase. Operators were blinded to treatment group for ultrasound and myography (below).

### Aortic myography and examination of side branches

Selected ApoE-/- male F1 adult offspring with and without maternal vape exposure underwent *ex vivo* aortic myography as previously described.^20,22,35^ The suprarenal aorta was explanted and transported in ice-cold PBS, and a 0.6-0.8 cm segment was secured on stainless-steel cannulas and mounted in the heated chamber of a pressure arteriography system (Model 110P, Danish Myotechnology, Denmark) and extended to *in vivo* length. Any side branches were surgically sutured. Physiological saline solution at 37°C, aerated with 5% CO2/95% O2 was used to fill the chamber and for aortic perfusion. After initial equilibration for 3 minutes, the aortic segments were pressured from 0 to 195 mmHg in 15 mmHg increments, and the outer diameter of the aorta segment was tracked by continuous computer video analysis measurement. The relative aorta outer diameter is presented as a ratio of outer diameter at baseline. For a subset of mice, side branches were visualized by injecting PBS containing 0.05% bromophenol blue into the apex of the heart at sacrifice after perfusion with PBS. The number of abdominal aortic side branches was recorded.

### Reduced representation bisulfite sequencing (RRBS) DNA methylation analysis

Aortic segments (suprarenal from ApoE-/- mice, infrarenal from C57BL6 mice) harvested from control (room air) F0 male and female mice, e-cig vaped F0 female mice, and male and female F1 offspring were subjected to RRBS Methyl-Seq (Active Motif, Carlsbad, CA) to identify differentially methylated genes and regions.

Genomic DNA (gDNA) from mouse abdominal aorta was extracted using Qiagen QIAmp DNA Mini Kit (Cat# 51304). The gDNA was quantified using Broad Range Reagents for the Qubit (ThermoFisher Scientific, cat number Q33231). Depending on DNA yield, samples were derived from either single mice, or two pooled mice per sample for both RRBS, and ATAC analysis (below). Fragment distribution was assessed on the TapeStation with gDNA ScreenTape and gDNA ScreenTape Buffer (Agilent, cat number 5067-5365). Samples that obtained satisfactory DIN score (>6) were used in the RRBS workflow. One hundred ng of gDNA was digested with MspI (NEB R0106) at 37⁰C overnight. Following enzymatic digestion, samples were used for library generation using the Zymo RRBS Library Kit (Zymo D5460) following manufacturer’s instructions. In brief, digested DNA was randomly ligated, and, following fragment end repair, bisulfite converted using Bisulfite Conversion component of the Zymo RRBS Library kit following the protocol. After conversion and clean-up, samples were amplified resuming the Zymo RRBS Library kit and Zymo-Seq UDI Primer Plate for library amplification and purification. Libraries were measured using Agilent TapeStation System and quantified using Qubit. Libraries were sequenced on an Illumina NGS platform using PE50 utilizing ∼45 million reads. This was followed by bioinformatic analysis using mm10 for alignment, mapping with Bismark v0.23.0, and differential analysis with methylKit 1.28.0. Significant thresholds: q-value <0.1, difference >0.25. Samples were evaluated by PCA, hierarchical clustering, and correlation.

Sites identified in this fashion were subjected to pathway analysis (utilizing QIAGEN IPA (QIAGEN Inc., https://digitalinsights.qiagen.com/IPA)^36^, DAVID^37,38^, and Enrichr^39^). Predicted/pseudo-genes were removed from the gene lists prior to performing analysis. For DAVID GO and KEGG enrichment, an EASE threshold of 0.05 was used with a “Mus musculus” background. For Enrichr Wikipathways (Mouse) an adjusted p-value of 0.05 was used. ChEA3 (ChIP-X Enrichment Analysis Version 3^40^) was used to identify enriched transcription factors predicted to target differentially expressed genes.

### Assay for Transposase-Accessible Chromatin (ATAC)-sequencing

Blood was collected from mice using cardiac puncture, and centrifuged at 2500 x g at 4 °C for 10 minutes. Plasma was removed and the cells were flash-frozen. Flash-frozen blood cell samples harvested from C57BL/6J control (room air) F0 male and female mice, e-cig vaped F0 female mice, and male and female F1 offspring underwent ATAC-seq using a kit (Active Motif, cat # 53150, Carlsbad, CA) to identify differentially accessible genes and regions.

The tissue was manually disassociated, isolated nuclei were quantified, and 50,000-100,000 nuclei were tagmented using Tn5 transposase and buffer provided in the ATAC-Seq Kit (Active Motif). Tagmented DNA was then purified using a DNA extraction method and amplified with PCR, adding a unique i5 and i7 index combination. The resultant libraries were purified using SPRI beads. The final libraries were quantified and sequenced using PE50 on the Illumina platform. Following FastQC generation, adapter trimming was performed using Trim Galore, alignment used the Burrows-Wheeler Aligner (BWA), followed by filtering out of duplicate reads, and mitochondrial DNA reads.

Only uniquely mapped reads (mapping quality ≥1) were used for further analysis. Alignments were extended in silico at their 3’-ends to a length of 200 bp and assigned to 32-nt bins along the genome. The resulting histograms (genomic “signal maps”) were stored in bigWig files. Peaks were identified using the MACS3 and Epic2 at a cutoff of p-value 1e-7, without control file, and with the –nomodel option. Peaks that were on the ENCODE blacklist of known false peaks were removed. Signal maps and peak locations were used as input data to Active Motif’s proprietary analysis program, which creates Excel tables containing detailed information on sample comparison, peak metrics, peak locations and gene annotations. Differential analysis was performed using DESeq2. Significance cutoff = adjusted p-value <0.05 and log-fold-change > 0.35. HOMER (Hypergeometric Optimization of Motif EnRichment) was used for motif analysis. Differentially accessible gene (DAG) pathway analysis was performed as previously described.^41^

### Statistics

Data are presented as mean ± SEM. Groups were compared using Student’s t-test (two-tailed) for parametric data. When comparing multiple groups, data were analyzed by ANOVA with Bonferroni’s post-test. Sequential measurements were analyzed by One-Way Repeated Measures ANOVA. Paired t-testing was performed utilizing Wilcoxon matched-pairs signed rank test with Spearman effectiveness testing. Chi-square testing with Fisher’s Exact test was used for ApoE-/- AAA incidence comparisons. Aortic diameters were compared using two-way ANOVA (α 0.05) with multiple comparisons. All statistic testing and graph composition was done using GraphPad Prism software version 11 (San Diego, USA). Proportionate Venn diagrams were created with DeepVenn (arXiv:2210.04597). A value of P ≤ 0.05 was considered statistically significant. The corresponding author had full access to all the data in the study and takes responsibility for its integrity and the data analysis.

## Results

### Maternal e-cig nicotine vaping augments offspring aortic stiffness and AAA development

Using the SciReq InExpose system, we performed one hour of daily vaping of adult female C57Bl/6J or ApoE-/- mice for 1 week, followed by mating to control males, and then continued daily vaping through gestation. The protocols used in these experiments achieved plasma cotinine levels similar to those seen in moderate human e-cig users and smokers (see previous publication^18^). A separate group of female mice were treated with vaping for 4 weeks using the same protocol, and then were harvested for DNA analysis (FVF0 = female vaped F0 generation - Subset A).

Offspring (FVF1/MVF1 = female/male maternally-vaped F1-generation) were aged to 10 weeks of age in room air, and then either tissues were harvested for DNA analysis (Subset B) or ex-vivo aortic myography, or the mice were subjected to AAA-induction (Subset C) via either continuous osmotic pump AngII-infusion (ApoE-/- mice) or infrarenal PPE infusion operation (C57Bl/6J mice). Offspring from room-air-treated mice were used as controls (males = MRA, female = FRA). See **Figure 1** for a schematic of the experimental workflow.

**Figure 1.**
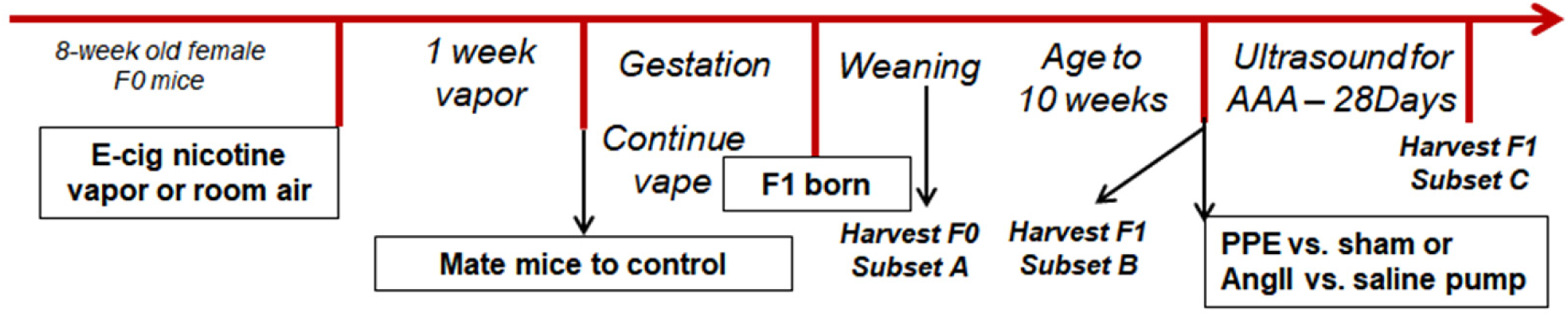
Schematic of Experimental Workflow. *Murine Group Acronym Definitions*: 1) FRA, MRA = F0 female and male mice (respectively), treated with room air 2) FVF0 = F0 female mice treated with vaping only, for 4 weeks (Subset A) 3) FVF1 = Adult female F1 offspring of vaped female mice (Subset B and C) 4) MVF1 = Adult male F1 offspring of vaped female mice (Subset B and C)

Increased aortic stiffness is thought to predispose to AAA development.^20,22^ Using ex-vivo arterial myographic studies of harvested suprarenal aorta from MVF1 ApoE-/- offspring mice (Subset B), we found significantly higher aortic stiffness than in MRA controls (**Figure 2**). A trend towards higher stiffness was also seen in FVF1 ApoE-/- mice (vs. FRA).

**Figure 2:**
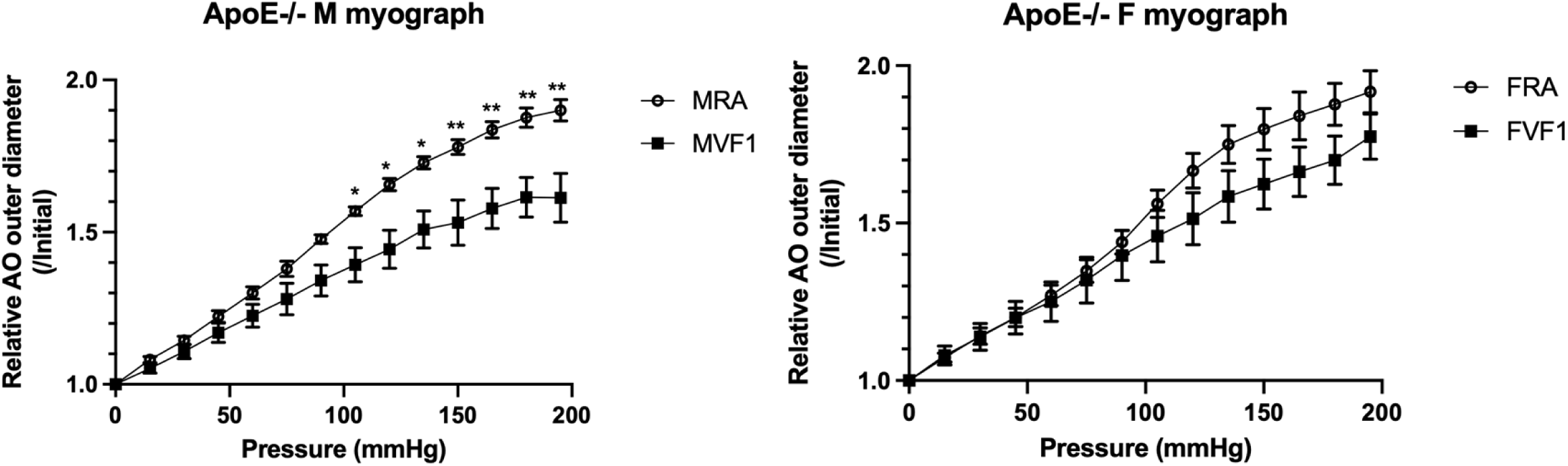
E-cig vaping in female F0 ApoE-/- mice augments aortic stiffness in their male F1 offspring. Ex vivo arterial pressure myography of supra-renal aortic segments. 8-week old ApoE -/- females were exposed to e-cig as described (or room air) which continued through mating and pregnancy. Offspring (MRA/FRA = male/female room air; MVF1/FVF1 = maternal e-cig) were aged to 10-weeks, and their explanted abdominal aortae were pressurized from 0-195 mmHg in 15 mmHg increments. Male (left, n = 5 both groups) and female (right, n = 4 both groups) aortic segments from animals with maternal e-cig exposure showed increased stiffness in response to increasing pressure levels, with significant augmentation in males, and up-trending in females. *p<0.05, **p<0.005 for specific pressure timepoints.

Intriguingly, anatomic studies showed an increase in the number of small suprarenal aortic side-branches in both MVF1 and FVF1 ApoE-/- and C57BL6/J mice (Subset B) when compared with maternal-room-air-exposed offspring, which was significant in ApoE-/- males and in both C57BL6/J males and females (**Supplemental Figure 1**).

AAA growth progression was serially tracked in Subset C mice by post-induction ultrasound. In both male and female offspring, and in both models, maternal e-cig exposure significantly augmented AAA development when compared with room-air (**Figure 3a-d, Supplemental Figure 10**). Growth vs. baseline was more pronounced in males in both models. Further, AngII-induced AAA incidence (>150% baseline diameter) in the room-air ApoE-/- mice (55% in males, 36% in females) was consistent with typical outcomes found in the literature, but significantly higher in MFV1 mice (90%), and with an upward trend in FVF1 mice (56%) (**Figure 3e**). No significant mortality differences were noted. Saline pump-infused (for ApoE-/-), and sham operated control (for C57BL6/J) mice showed no significant growth in any group (data not shown).

**Figure 3A-D:**
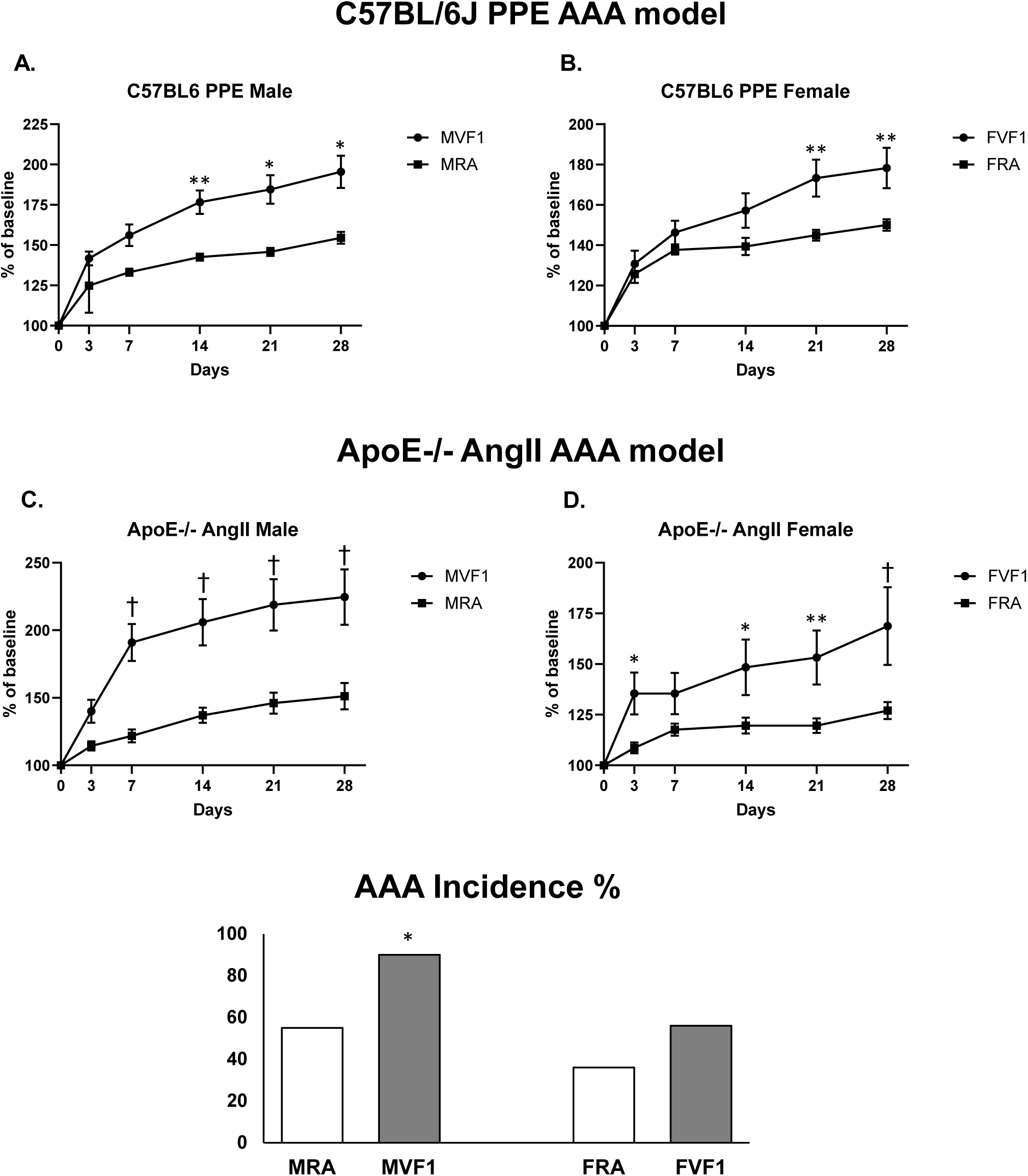
F1 Generation AAA Growth Curves. **A/B.** Male and female C57BL/6J mice, PPE – porcine pancreatic elastase-induced infrarenal AAA; **C/D.** Male and female ApoE-/- mice, AngII – angiotensin II pump-induced suprarenal AAA. AAA progression in F1 offspring, with maternal vaping (MVF1/FVF1) vs. maternal room air exposure (MRA/FRA). **A.** Left, males (n = 10 MVF1, 11 MRA). **B.** Right, females (n = 12 FVF1, 10 FRA). **C.** Left, males (n = 10 MVF1, 30 MRA). **D.** Right, females (n = 9 FVF1, 26 FRA). Standard error shown. M/FVF1 aortic diameters grew significantly larger vs. room air (mixed-effects analysis, Bonferroni repeated measures multiple comparisons test), *p<0.05, **p<0.005, †p<0.0001. **Figure 3E. Incidence of AngII-induced AAA (>150% of baseline size) in ApoE-/- mice.** Room air, n = 33 males (MRA), 25 females (FRA); MVF1 = males with vaped mothers, n = 10; FVF1 = females with vaped mothers, n = 9. *p<0.05 vs. room air.

### E-cig vaping alters aortic DNA methylation in female mice

We performed RRBS analysis in the C57BL/6J mice (Subset A) to examine DNA methylation changes which occurred in infrarenal aortic tissue in female mice after 4 weeks of e-cig nicotine exposure (vs. room-air). The final set consisted of two FRA and three FVF0 samples, which clustered independently when subjected to hierarchical clustering and principal components analysis (**Supplemental Fig 2**).

We found that e-cig exposure dramatically and significantly altered the aortic epigenome, in thousands of CpG locations. We identified 51,012 differential bases in 2,390 regions with a q-value threshold <0.1 and difference threshold >0.25, involving a total of 9,824 Differentially Methylated Genes (DMGs) of which 3,508 were hypomethylated with vaping and 8,770 were hypermethylated (some genes displayed both processes, but in different CpG locations). Among those were 4,772 differentially methylated promoters, 3,245 CpG islands, and 7,624 genes with intron or exon methylation differences (**Supplemental Table 1a**).

Pathway analysis of the involved aortic tissue genes identified changes in numerous transcription factors, cellular process pathways (e.g. differentiation, migration, proliferation, apoptosis, adhesion), and a number of immune/inflammatory-related processes including the IL6, TGF-beta, MAPK, and NF-kappa B signaling pathways as well as T-cell differentiation (including Th17, Th1 and Th2 cells) and leukocyte trans-endothelial migration. Also noted were many developmental and homeostasis-related “-signaling” pathways (e.g., Hippo, Rap1, MAPK, mTOR, apelin, Wnt, calcium, Notch, BMP, PI3-Akt), and others related to smooth muscle function, matrix remodeling, and atherosclerosis (**Figure 4**). Analysis also revealed major changes in pathways related to the terms “blood vessel development” “angiogenesis”, and “vasculogenesis”, suggesting vape exposure potentially leads to dysregulation of dozens of these genes in the aorta after a 4-week exposure (**Supplemental Table 1d**). In general, metabolic pathways and SMC contractile gene loci tended to be more significantly hypermethylated, while immune pathway gene loci were more significantly hypomethylated – suggesting predisposition to a pro-inflammatory state accompanied by SMC phenotypic switching and de-differentiation.

**Figure 4:**
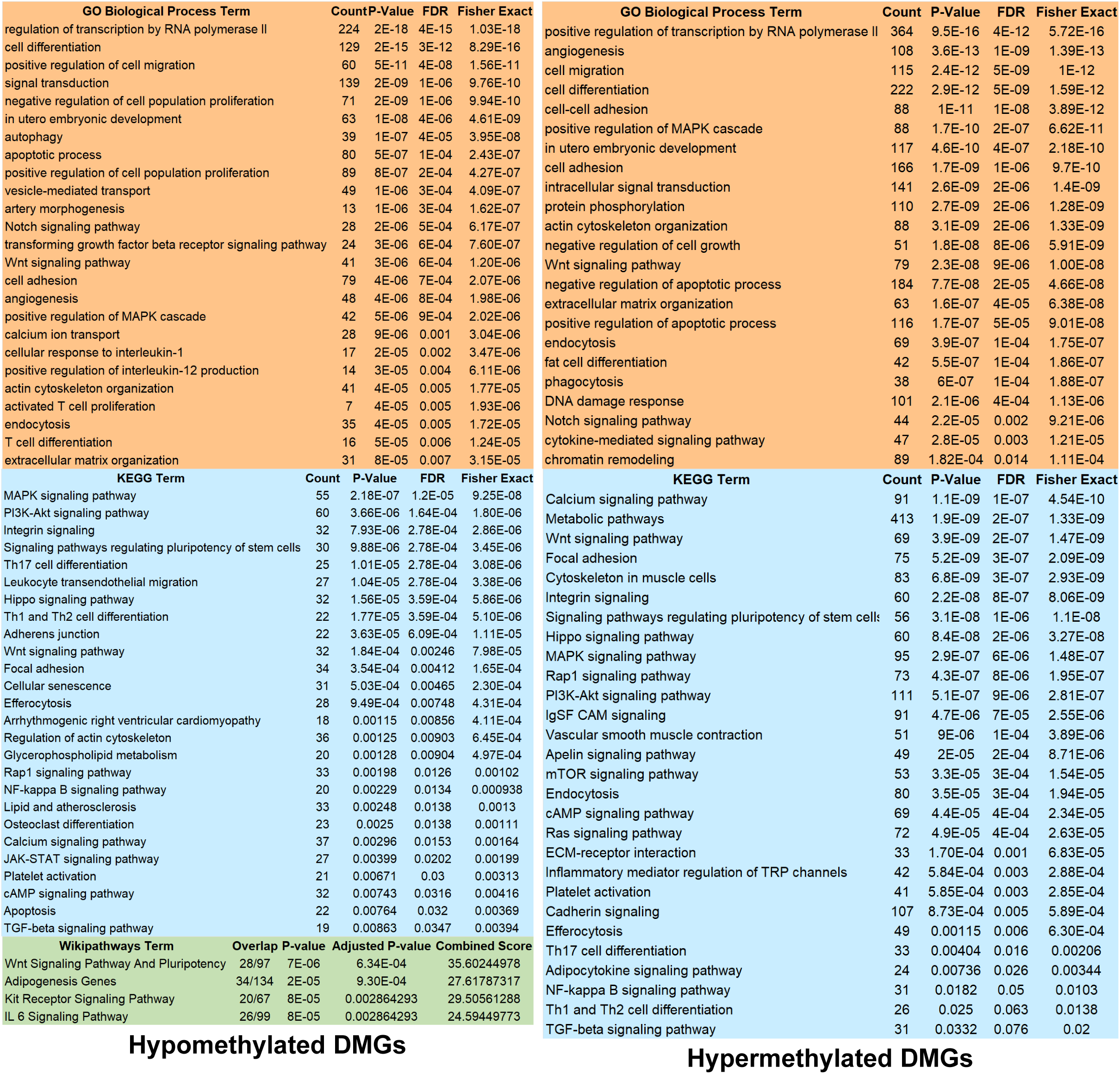
Pathway analysis of FRA (female control) vs. FVF0 (e-cig treated female) DMGs (C57BL/6J mice). Gene Ontology (GO) Biological Process, KEGG, and Wikipathways: analysis performed as described in Methods. Selected vascular-tissue-relevant pathways shown. Hypo = less/Hyper = more CpG methylation with treatment (vs. control). Left – hypomethylated DMGs; Right – hypermethylated. Count = number of DMGs in pathway, Overlap = number of DMGs in pathway over total genes in pathway. Various enrichment statistics shown. Wikipathways combined score derived by multiplying unadjusted Fisher’s exact test p-value by Z-score deviation from the expected rank. Hypermethylated DMGs did not yield significant Enrichr Wikipathways. Complete pathway analyses with genes in each category are found in Supplemental Tables.

Notably, when the list of significant DMGs was compared with a list of 120 prioritized human genes containing AAA-risk-related variant loci from our recent co-authored AAA GWAS study^42^, there were 42 overlapping genes – suggesting that vape exposure in mice rapidly leads to epigenetic aortic DNA remodeling in a manner that may predispose to AAA development. (**Supplemental Table 2**). These included genes such as CHRNA5, PLAU, LDLR, RORA, and SPP1. These results complement our previous work which demonstrated that exposure to e-cig vapor augmented AAA development in two different models.^18^

### Maternal e-cig vaping leads to durable epigenetic changes in offspring

We sought to clarify the mechanisms behind the augmented AAA development observed in offspring of vape-exposed mice. Therefore, we performed aortic tissue high-throughput RRBS DNA methylation analysis of Subset B (10-week-old offspring of vape-exposed mice), and compared them with same-sex room-air-parent control mice. Offspring samples were products of different litters for this analysis, and for the ATAC-seq analysis described below. Similar to the findings described above, we found numerous vape-induced CpG methylation changes, in thousands of gene loci.

Comparing female (FVF1) Subset B samples with control (FRA) (n = 2 in each) we identified 22,257 differential bases in 614 regions (thresholds: q-value <0.1, difference >0.25) comprising 6,448 combined DMGs. As before, the preponderance of differentially methylated CpG changes consisted of hypermethylation (1,833 with hypomethylated CpGs, and 5,604 with hypermethylated CpGs) (**Supplemental Table 1b**) with clear differential clustering (**Supplemental Figure 3**). Differential bases were found in the promoters of 1,758 genes, in the introns/exons of 4,697 genes, and in CpG islands near 373 genes. Notably 5,635 of the F1 generation female DMGs (87.4%) overlapped with those from the FRA vs. FVF0 (Subset A) list described above, suggesting a durable cross-generational e-cig-related epigenetic signature in vascular tissue (**Figure 5**). Consistent with this finding, pathway analysis of the F1 female DMGs identified largely similar pathways to those seen with direct maternal exposure (**Figure 6a**).

**Figure 5:**
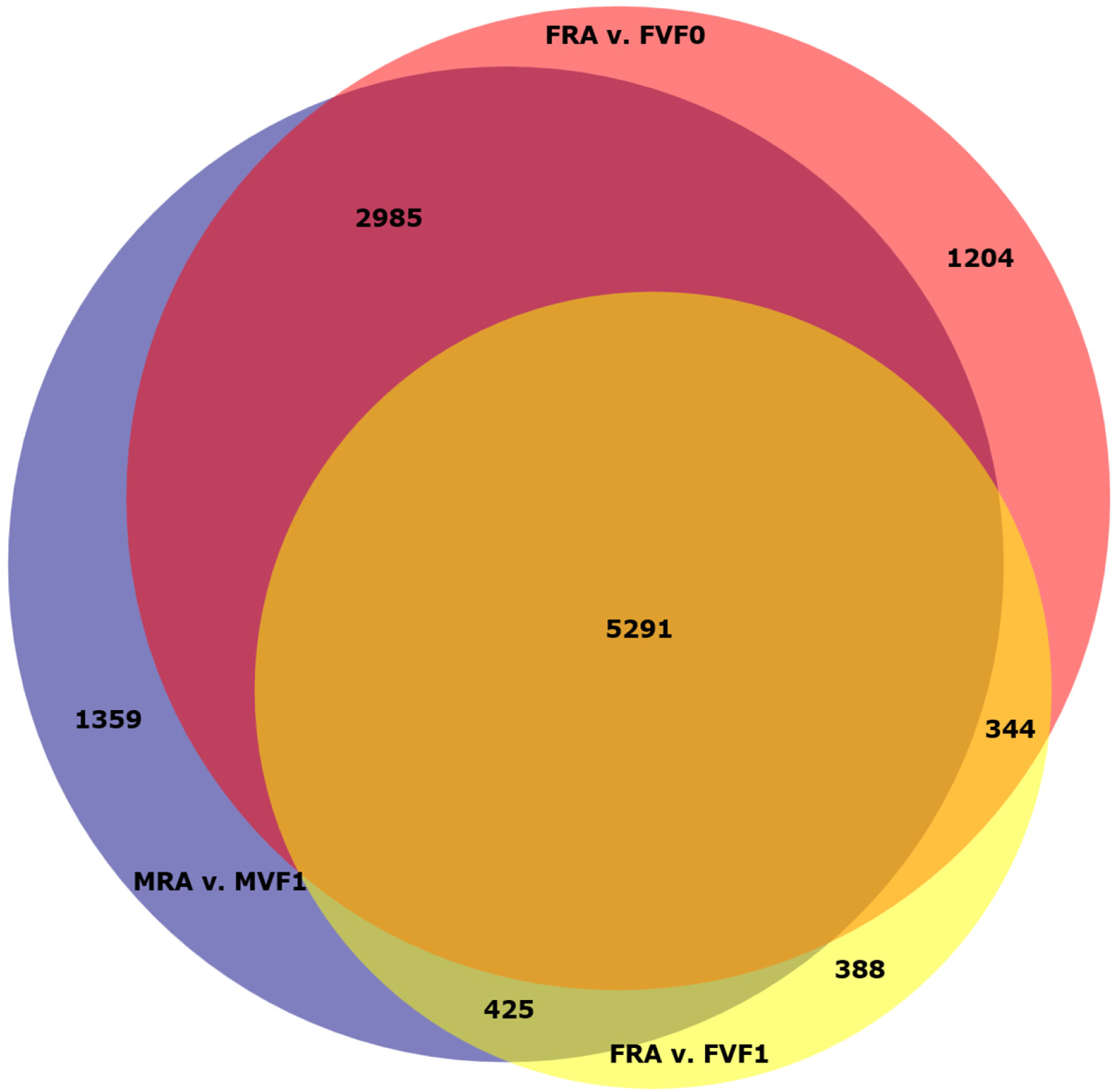
Female-Male Aorta RRBS DMGs Venn. Proportionate Venn diagram showing intersecting gene lists of significant DMGs (C57BL/6J mice). Yellow = FRA vs. FVF0 (Subset A), Red = FRA vs. FVF1 (Subset B), Blue = MRA vs. MVF1 (Subset B).

**Figure 6A-B:**
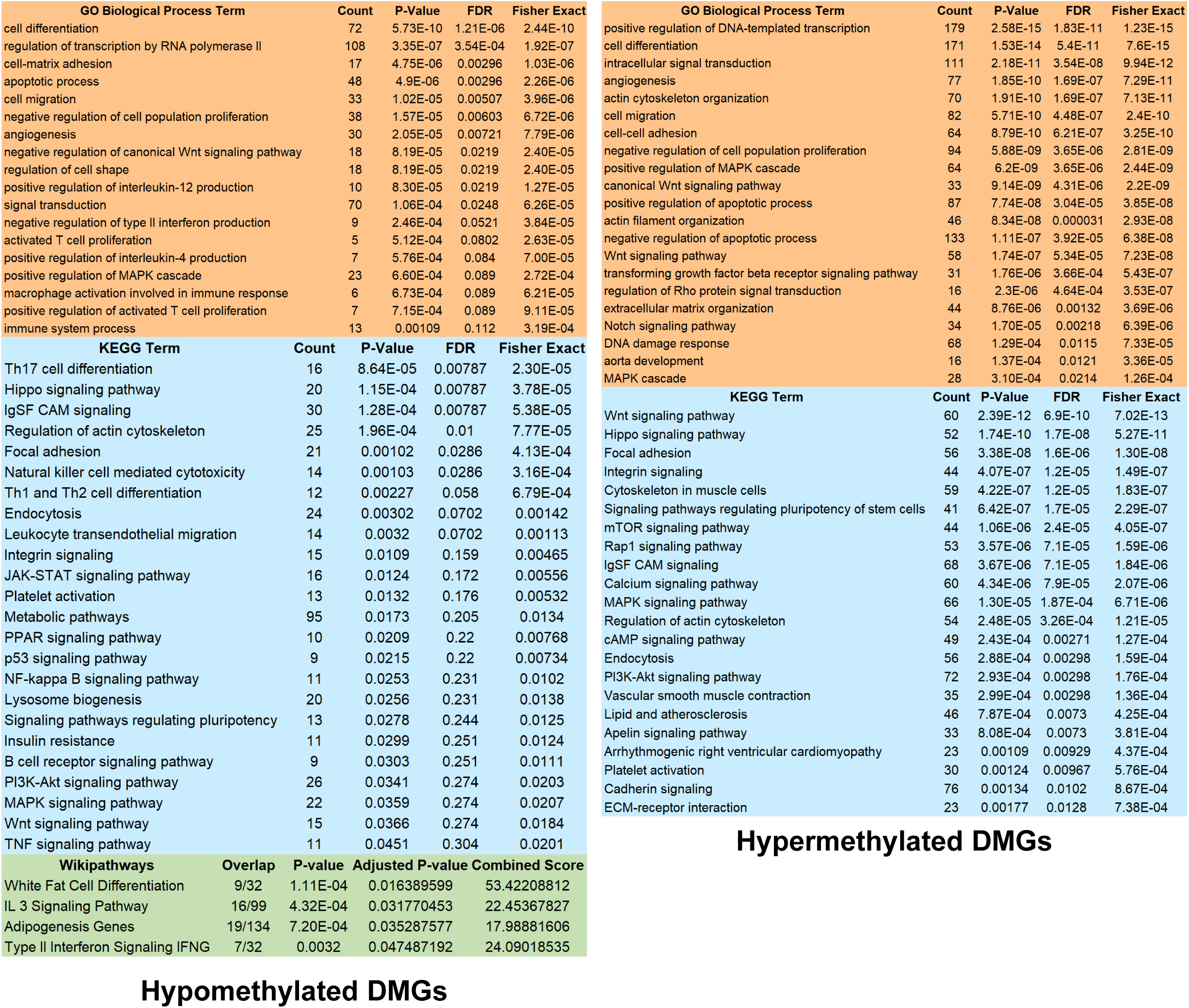
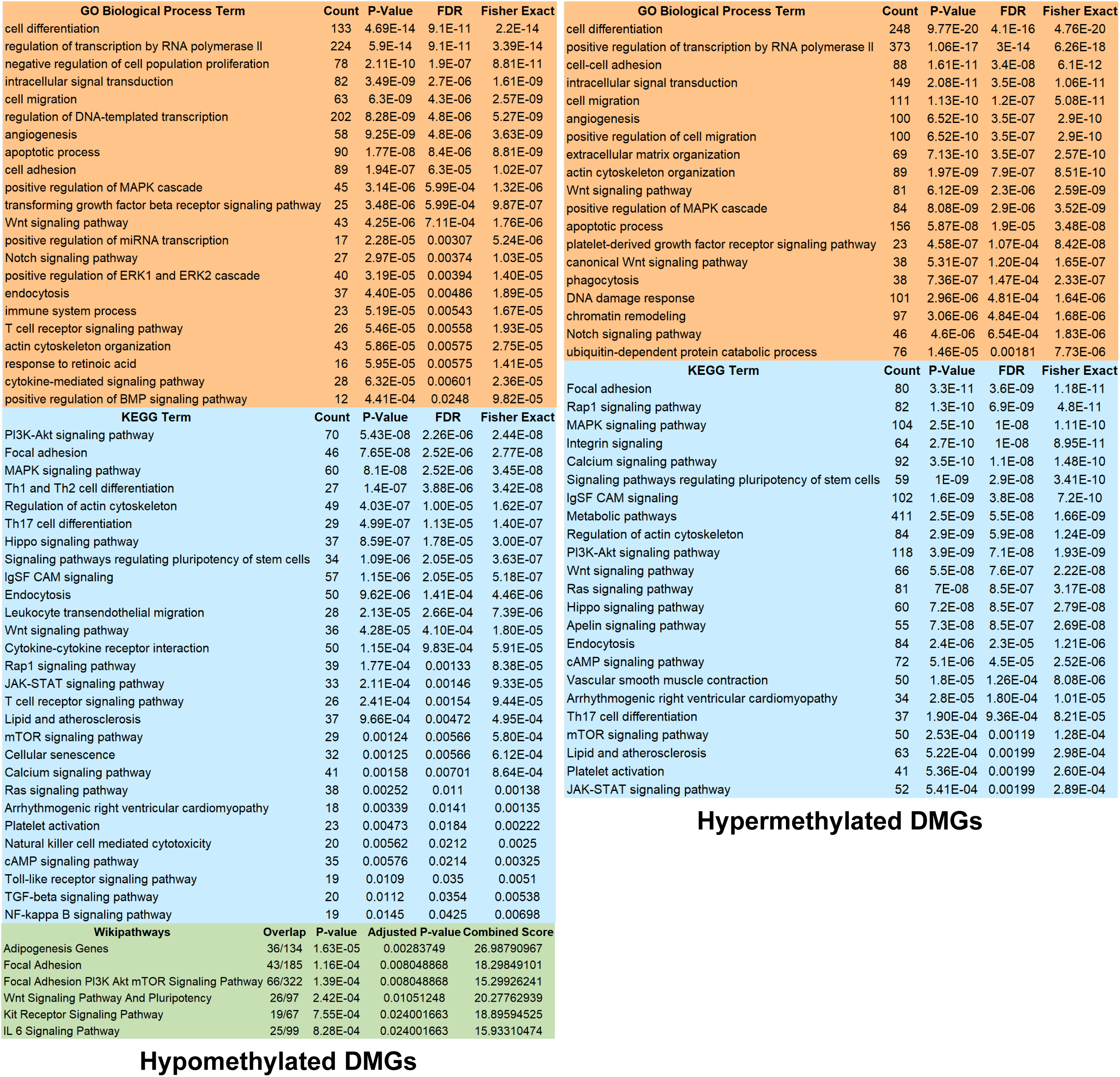
F1 Generation DMG Pathway analysis. A (above). FRA (female control) vs. FVF1 (female offspring of e-cig treated females. **B (below).** MRA (male control) vs. MVF1 (male offspring of e-cig treated females). C57BL/6J mice. Gene Ontology (GO) Biological Process, KEGG, and Wikipathways: analysis performed as described in Methods. Selected vascular-tissue-relevant pathways shown. Hypo = less/Hyper = more CpG methylation with treatment (vs. control). Left – hypomethylated DMGs; Right – hypermethylated. Count = number of DMGs in pathway, Overlap = number of DMGs in pathway over total genes in pathway. Various enrichment statistics shown. Combined score derived by multiplying unadjusted Fisher’s exact test p-value by Z-score deviation from the expected rank. Complete pathway analyses are found in Supplemental Tables.

We also examined male offspring Subset B aortic tissue (n = 3) using the same methods and compared them with room-air exposed males (n = 3), identifying considerably more DMGs than were seen in females. As noted for both other comparisons, the preponderance of changes displayed hypermethylation.

We identified 59,503 differential bases in 2,738 regions, comprising 10,060 combined DMGs using the above thresholds (3,903 DMGs with hypomethylated CpGs, and 8,922 with hypermethylated CpGs) (**Supplemental Table 1c**). Clustering again showed clear group separation (**Supplemental Figure 4**). Differential bases were found in the promoters of 3,378 genes, in the introns/exons of 7,330 genes, and in CpG islands near 677 genes. A substantial number of genes (8,276 genes) from the male analysis (82.3%) overlapped with the maternal Subset A analysis, again suggesting a consistent and durable cross-generational aortic methylation signature. Further, a remarkable 5,291 DMGs were found to overlap between all three RRBS analyses (**Figure 5**). As expected from the level of overlap, pathway analysis of the male DMGs strongly resembled that of the other two described comparisons (**Figure 6b**).

Genes associated with murine AAA development were obtained from our previous publication (see NCBI GEO GSE51228)^43^, in which we performed whole-genome microarray transcriptional profiling of infrarenal aorta 7 days after PPE-induction (scr-miR-treated PPE vs. sham-operated control). The ChEA3 online tool^40^ was used to identify the enriched transcription factors predicted as most likely to target the genes that were significantly up- or down-regulated with AAA from the published dataset. These were then compared with the list of differentially methylated transcription factors (TFs) derived from the aortic DMGs from this study which, as noted above, were prevalent. Of the 100 highest-ranked (“Mean-Rank”) enriched transcription factors targeting AAA up- and down- regulated genes, 35 and 36 respectively were found to be aortic DMGs in the female Subset A comparison (**Supplemental Table 4**). Further, 28 and 30 TFs overlapped respectively with female Subset B DMGs, and 35 and 32 TFs with male Subset B DMGs. These findings were remarkably consistent between the sets – a total of 26 and 27 (respectively) of the TFs were found to be in-common between all three DMG sets. These data strongly imply that vape-treatment differentially methylates transcription factor DNA in a fashion that predisposes to activation (or suppression) of AAA-related genes in aortic tissue, and that these changes remain durable to adulthood in their offspring.

### E-cig vaping alters DNA accessibility in blood cells of female mice and offspring

Blood was harvested from parallel treated and control C57BL/6J F0-generation female mice and their F1 progeny (Subsets A and B) and the cellular DNA was isolated and evaluated with ATAC-seq to examine the impact of e-cig nicotine vaping on DNA accessibility (adjusted p-value <0.05). The results revealed numerous differentially accessible genes (DAGs) with (vs. without) e-cig exposure, both in the F0 mothers and their F1 offspring (**Supplemental Tables 3a-c**).

Comparing Subset A females (room-air treated females (FRA) vs. vaped females (FVF0), n = 3 per group), we found treatment led to 1,038 more accessible vs. control (“UP”) peaks, and 802 less accessible (“DOWN”) peak regions (totaling 784 and 721 DAGs respectively, with 12 overlaps) (**Figure 7**). DAG UP pathway analysis showed enrichment for numerous transcription factors, and AAA-related pathways such as T-cell and B-cell differentiation, immune and inflammatory responses (including TNF and interferon signaling), TGF-beta signaling, apoptosis, MAPK signaling, and chromatin remodeling, as well as EGFR1 signaling and genes related to adipogenesis and insulin signaling (see **Supplemental Table 5** for full analysis results). Enriched DAG DOWN pathways included transcription regulators, cell proliferation and migration, and blood vessel morphogenesis- angiogenesis genes. These results mirror those seen with aortic DNA methylation analysis for these same mice.

**Figure 7.**
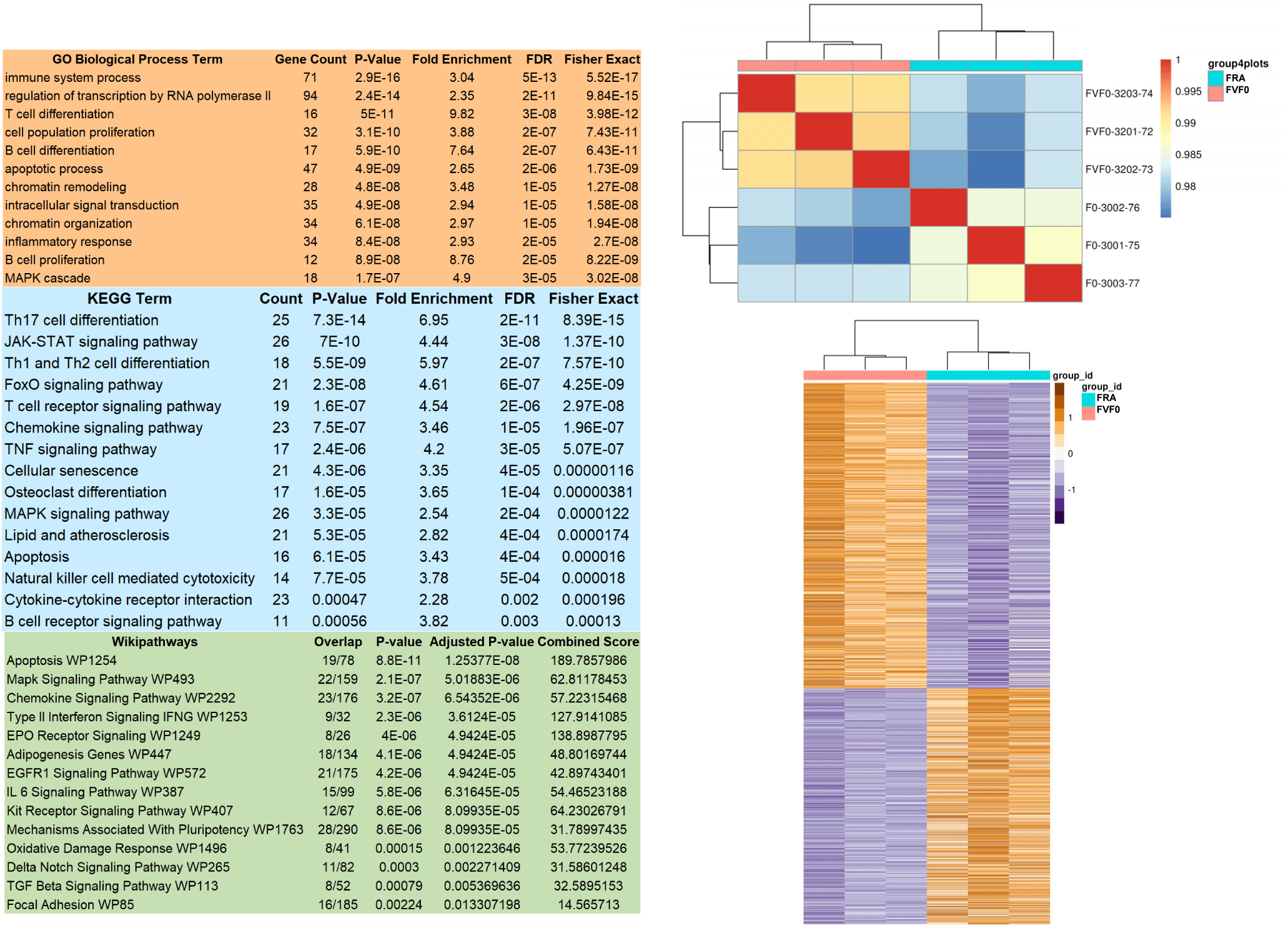
Direct impact of vaping on F0 (FRA) C57BL/6J female mice: Blood ATAC-Seq. Analysis identified numerous differential accessibility genes (DAGs, pAdj<0.05, logFC >0.35) in response to 4 weeks of vaping. Read range: ∼86-151 million; FRIP score range: 14-43%. DAGs vaped vs. room air (784 UP with vape). Left, Gene Ontology (GO) Biological Process, KEGG, and Wikipathways, analysis performed as described in Methods. Selected vape-enhanced pathways shown. Top right, sample PCA Pearson Correlation; Bottom right, heatmap of all significantly regulated DAGs (blue = room air controls, pink = e-cig vaped).

Ingenuity pathway analysis (IPA) was also used to probe the Subset A comparison UP DAGs and construct regulator effects networks. Notably, both the #1 and #2 highest consistency rank networks predicted the substance nicotine to be an upstream regulator (**Supplemental Figure 5a**), providing validation that nicotine was the primary effector for the observed epigenetic alterations. Similar to the findings shown using other methods, IPA canonical pathway enrichment of more accessible loci was dominated by immune system activation and cytokine signaling (**Supplemental Figure 5b**). An IPA summary graph network identified key inflammation-related hubs including CD28, IL1b, IFNG, TNF, STAT1, AKT1 and SOCS1 (**Supplemental Figure 5c**). These results are consistent with our prior work suggesting that nicotine e-cig vaping is associated with augmented inflammatory signaling.^18^

The top 3 ranked HOMER analysis de novo motifs for the UP peaks included ELK3, IRF4 and IKZF1, with the HOMER Known motif analysis noting many ETS family and IRF family motifs. For DOWN peaks, de novo significant motifs included BORIS/CTCFL, Sp1 and PU.1, while the highest ranked HOMER Known motifs primarily featured zinc fingers such as CTCF, BORIS, KLF1/3/4/5 and Sp1/2/5 (**Supplemental Figure 6** and **Supplemental Table 6**).

Looking at female (FVF1) Subset B samples against control (FRA) (n = 3 per group) yielded 9,504 peak regions, associated with 2,857 UP (more accessible vs. control) DAGs and 2,493 DOWN DAGs (with 221 genes on both lists) (**Supplemental Figure 7**). When comparing the more accessible DAGs (UP) between the female Subset A (F0) and Subset B comparison (F1), we observed 77.6% (608 genes) overlap between the gene lists (**Figure 8a**). There was also 58.3% overlap (421 genes) in the less accessible DAGs (**Figure 8b**). UP pathway enrichment revealed generally the same pathways as for Subset A, with clear immune/inflammatory signaling and transcriptional regulators prominently featured (**Supplemental Table 7**). DOWN pathways were again enriched for transcription factors, angiogenesis and cell proliferation and migration genes, as well as cytoskeletal genes and extracellular matrix organization, and also included developmental and differentiation pathways such as canonical Wnt signaling and Hippo signaling.

**Figure 8A-B:**
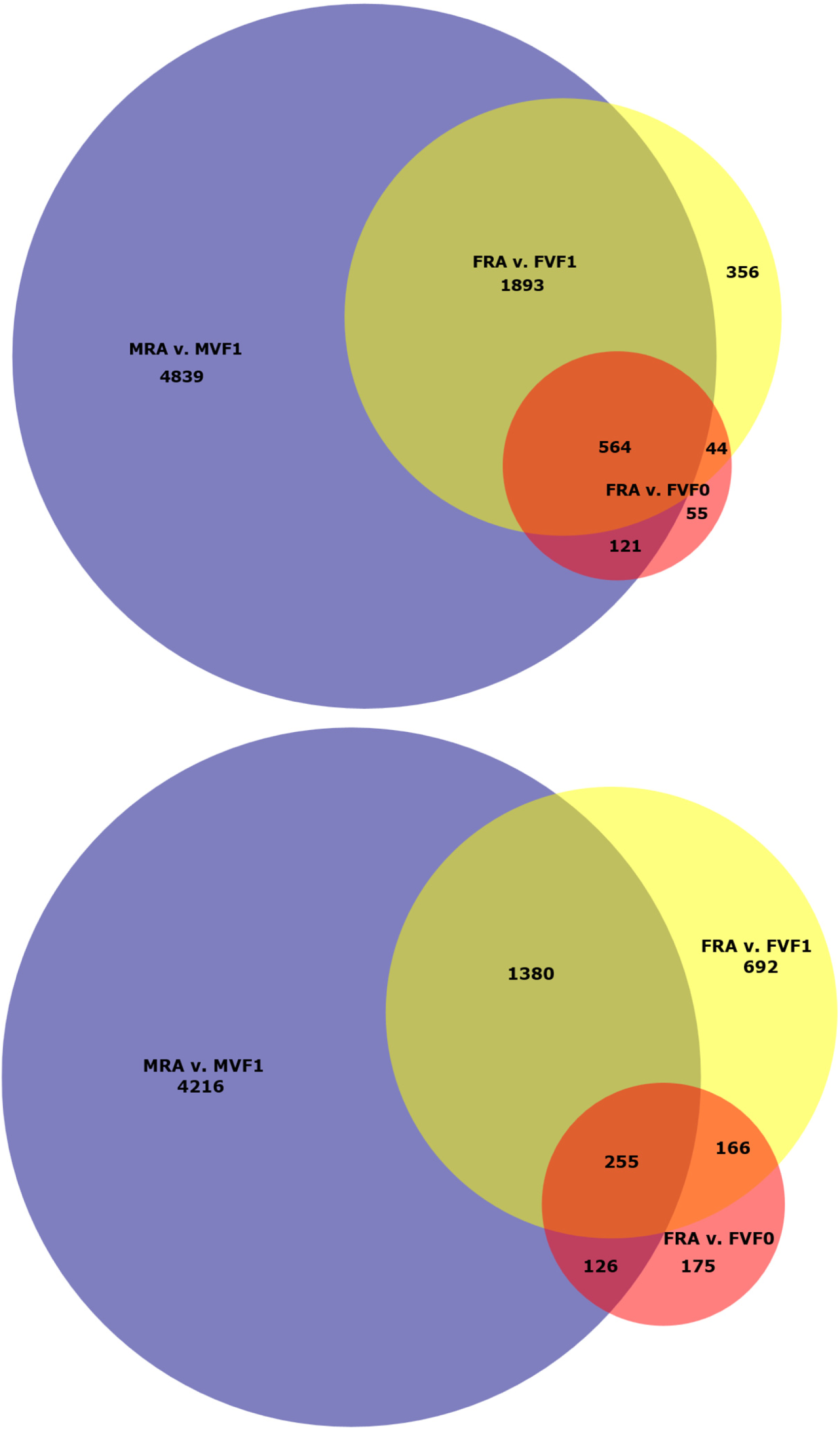
ATAC DAG Venn Diagrams. A (top), Female-Male Blood UP DAGs. **B (bottom),** Female- Male Blood DOWN DAGs. Proportionate Venn diagrams showing intersecting gene lists of significant DAGs (C57BL/6J mice). UP = more accessible, DOWN = less accessible, with treatment. Yellow = FRA vs. FVF0 (Subset A), Red = FRA vs. FVF1 (Subset B), Blue = MRA vs. MVF1 (Subset B).

HOMER Known and de novo motif analysis of the female F1 generation DOWN peaks showed similar findings to that seen above with the F0 generation, with top ranked TFs including BORIS/CTCFL, but also HIC1/Treg-ZBTB29, and with zinc fingers (like CTCF, BORIS and Sp1/2, but also Gata2/4/6) and basic leucine zipper (bZIP) motifs (e.g. HLF, NFIL3). As before, UP motifs featured primarily ETS and IRF family factors and some Runt family, with top-ranked de novo motifs including ETS1, PU.1-IRF, and RUNX2 (**Supplemental Figure 8** and **Supplemental Table 8**).

Male sample comparison (MVF1 (n = 3) vs. room-air males (n = 2)) yielded considerably more peaks and DAGs than in their female counterparts, including 23,315 significant peak regions, representing 7,417 UP (more accessible than in controls males) and 5,977 DOWN DAGs (with 2,296 overlaps) (**Supplemental Figure 7**). Comparing the more accessible (“UP”) DAGs for the male vs. female (F1), we found that 2,457 genes (86% of the female list) appeared on both lists, while 1,635 DOWN DAGs overlapped (65.6% of the female list) (**Figures 8a-b**).

Unsurprisingly, given the considerably larger number of genes identified in the male group, pathway analysis yielded many more significant categories (**Supplemental Table 9**). However, overall the pathway types seen were the same ones enriched in the female Subset A and Subset B groups. For UP DAGs we again observed major representation of chromatin remodeling and transcriptional regulation in the enriched pathways, along with the previously noted apoptosis, T-cell/B-cell and numerous immune/inflammatory pathways. More prominently featured were pathways related to DNA damage/repair responses and protein ubiquitination, as well as Notch signaling. DOWN DAGs also showed the same pathway categories as for the other comparisons, although they included some inflammatory pathways as well, possibly due to many inflammatory/immune genes undergoing complex regulation. As an example, of 154 genes in the “inflammatory response” GO Biological Process category from the DOWN DAGs, 57 of them also were found in the UP DAG list.

HOMER motif analysis was more complex for the male comparison, with highly-ranked UP de novo motifs including Fli1, Sp2, STB5, NFY, NRF1 and RUNX, and known motifs again featuring ETS, IRFs, Runts and zinc fingers. Top-ranked DOWN motifs included SpiB, FOS/JUNB, and KLF5, with most known motifs originating from the bZIP (e.g. Fos, JunB, Atf3, Fra1) and ETS (e.g. PU.1, Elf4/5, ETS1) families (**Supplemental Figure 9** and **Supplemental Table 10**).

Promoter hypomethylation and gene body hypermethylation are both thought to generally correspond to activation, as does increased ATAC-detected DNA accessibility. Accordingly, we collected the DAG/DMGs from all male and female Subsets sharing those characteristics and looked at in-common genes, yielding a total of 96. Similar to the larger-scale analyses detailed above, DAVID pathway analysis identified these primarily as transcription factors (e.g. Irf2/Irf4, Lef1, Trp53, Klf2/3, Arid1b, Bach2, Smad7 and Egr2), immune system response genes (particularly T-cell/Th17/Natural Killer cell differentiation and Type 2 interferon signaling genes), chromatin remodeling genes (e.g. Ncoa3, Setd1b, Jarid2, Mta3), Wnt signaling, MAPK cascade (e.g. Dusp2, Map3k5/4k4) and lipid/atherosclerosis genes (e.g. Abcg1, Arhgef1, Ppp3ca).

Taken together, these results show that murine maternal e-cig exposure leads to changes in numerous epigenetic loci in blood and aortic tissue in both the mother and her offspring, many of which were conserved across generations and between genders, remaining durable to adulthood in the F1 generation. The observed changes suggest potential activation (or priming for activation) of processes known to be involved in both human and mouse-model AAA.

## Discussion

AAA rupture carries a mortality of 80%.^2^ AAA prevalence remains high in recent reports^44^, and although improved endovascular and open surgical techniques have led to reduced mortality^45^, deaths due to ruptured AAA and its complications are still common.^2^

Smoking is associated with an approximately 5-fold increased risk of AAA.^46^ Indeed, AAA is more closely associated with cigarette smoking than with any other tobacco-related disease except lung cancer.^47^ While smoking prevalence has generally declined in the U.S. population, e-cig use has exploded. One 2018 study estimated there were 10.8 million adult e-cig users in the U.S., of which 51.2% were under age 35^48^, and 2023 CDC data suggested that 6.5% of adults in the U.S. over age 18 were current e-cig users, including 15.5% of all adults aged 21-24. While men use them more often, 5.5% of all adult women also employ them (NCHS Data Brief No. 524, January 2025). As noted above, e-cig use during pregnancy is also prevalent. While the health effects of inhaling e-cig vapor are not yet fully known, vaping increases circulating levels of cotinine, the main nicotine metabolite, to or above levels reached with tobacco cigarettes.^49^ Further, multiple studies have suggested that e-cig vaping can have cardiovascular consequences,^10–14^ and does have significant short and medium term health impacts on children and young people.^50^

Despite the dire medical need, there is a relative paucity of published data exploring the role of nicotine in general, and e-cig vaping in particular, in AAA development. Buckley et al.^51^ found increases in AAA growth in 129/SvEv mice exposed to cigarette smoke. We have used two complementary murine models to show that nicotine infusion alone dramatically enhances AAA development.^17^ Further, we and our collaborators have found that both chronic infusion of nicotine, and nicotine-containing e-cig exposure induce murine aortic remodeling and stiffness, elements predisposing to AAA.^22,23^ Lastly, we have shown that e-cig nicotine vaping exposure augments aortic inflammation and AAA development in murine models.^18^

Convincing data also suggest that nicotine exposure has potential transgenerational consequences. Perinatal maternal nicotine in rats induced asthma in F1 and F2 offspring, with durable changes in global DNA methylation and histone acetylation in ovary, testis and lung.^52^ Pre- and perinatal maternal nicotine also affects the cardiovascular system in offspring. In rats it led to altered vascular contractility, heightened oxidative stress, increased blood pressure and aortic physiologic responsiveness to angiotensin II, as well as increased cardiac ischemia sensitivity in adulthood.^53–56^

In this study, we now show that maternal exposure to nicotine e-cig vapor significantly augments AAA in their adult male and female offspring, in two different strains, and using two different aneurysm models. While these findings are consistent with our prior data showing that nicotine vaping directly increases murine aneurysm growth^18^, it is crucial to highlight that we now also observe such effects after *in utero* exposure; this despite months of non-exposure after gestation.

As mentioned, the effects were most prominent in males, which is commonly observed in murine AAA models. However, it is likewise notable that females showed significant findings, as they have often been excluded from prior abdominal aneurysm-model studies, in part due to less impressive growth metrics. Sex-specific differences in environmental toxin processing, including sex-chromosome-dependent gene regulation and varying hormonal responses to inflammation and oxidative stress, might explain some of the observed differences. Indeed, females are believed to display more robust protective responses to genotoxic stress and physiologic DNA damage.^57^

Exposure to nicotine vaping in C57BL/6J mice led to numerous and widespread alterations in DNA methylation and DNA accessibility in abdominal aorta and blood, respectively. These changes were, to a large extent, conserved from parent to adult offspring, suggesting durability. As described in detail above, the observed changes suggest possible priming for activation of AAA-associated genes and pathways. Nicotine vaping especially seems to epigenetically alter loci associated with transcription factors, particularly ones known or predicted to regulate AAA-related pathways and processes.

Intriguingly, many intergenerational epigenetic alterations were found in angiogenesis- vasculogenesis genes. Nicotine treatment has been linked to altered vessel extracellular matrix remodeling, a key element in AAA formation.^58,59^ In the ApoE-/- mice *in utero* exposure appeared to have developmental impacts on the abdominal aorta, leading to increased aortic stiffness in offspring as adults. Further, both mouse strains that we used showed an increase in side-branch development in the supra-renal region in offspring. It has been suggested that aortic growth in the ApoE-/- angiotensin II AAA model relies in part on small side-branch vessel dissection events.^60^ The above-described changes might therefore have contributed in part to augmentation of AAA in this model. However no differences were seen in infra-renal branch numbers in the C57BL/6J mice (data not shown), which is the aneurysm location for the PPE model.

The amounts of vaporized nicotine utilized in this study raised murine cotinine levels to those comparable with moderate-to-heavy human use. However, e-cig vapor also typically includes aerosolized carrier fluid (most commonly propylene glycol and vegetable glycerol) as well as acrolein and formaldehyde. Our recent publication suggests that chemical constituents of e-cig aerosol may alter endothelial activation markers, and extracellular remodeling proteins.^23^ Exposure to these and other vaporized substances, such as heavy metals, may also have an impact on epigenomic re-programming. Notably, however, the DNA accessibility changes seen in the F0 generation females were predicted to have arisen primarily from nicotine as an upstream effector.

Genome-independent transcriptional regulation is achieved through epigenetic mechanisms, such as chemical modifications to specific regions of DNA (e.g. methylation) and their associated proteins (e.g. histone modifications, chromatin remodeling) that can silence or activate genes.^61^ These processes are complex, and regulate nearly all biologic pathways. DNA methylation (addition of a methyl group to 5’-cytosines in CpG dinucleotides) is performed by DNA methyltransferase (DNMT) enzymes, and largely manifests as a quantitative trait, altering relative gene expression. In general, genes whose promoters are methylated tend to be silenced, while “in-body” gene modifications are often activating.^26^ We found that e-cig exposure led to changes in chromatin remodeling and DNA damage-repair genes in parents and offspring, which could potentiate long-term impacts.

There have been numerous publications evaluating the impact of smoking and nicotine exposure on DNA methylation in humans.^27,28,62–66^ Cigarette smoke is among the strongest environmental modulators of DNA methylation.^25^ Nicotine in smoke downregulates DNMT1 (the primary DNA methylation repair enzyme)^24^ and can pass through the placenta to affect the fetus.^67^ Prenatal smoke exposure alters DNA methylation in offspring which can persist into adulthood.^26^

A recent study showed that e-cig use induced numerous carcinogenesis-related changes in DNA methylation in oral epithelial cells, with patterns similar to those seen in cigarette smokers^28^. Also, maternal tobacco use alters DNA methylation at imprinted genes, particularly affecting imprinting control regions and nearby enhancers in both the placenta and the developing fetus.^64^

It is unclear to what extent our observed augmented AAA growth and epigenomic changes in the mouse offspring were caused by direct trans-placental exposure to nicotine and vape by-products *in utero*. Other probable mechanisms include alteration of the uterine environment, or direct chromatin remodeling impacts on maternal germ-cell DNA prior to conception. Notably we found that several trans-generationally conserved DMGs were found to either be maternally imprinted genes, or factors involved in imprinting, such as H13, Gnas, Smoc2, Kcnq1, Dnmt3a, Arid4a, Kdm1b, and Mecp2. DNA methylation modifications in imprinted genes frequently lead to altered transcription and development, and can escape reprogramming during germ cell development, thereby resulting in durable alterations.

Huang et al. utilized an epigenome-wide association study (EWAS) to examine the association between urinary total nicotine equivalents and differential CpG loci in blood.^65^ Of the 230 human genes with differential CpGs with murine equivalents that they identified, 75 were found to be aortic DMGs in our e-cig vaped F0 females. Remarkably, we found that a large number of these same DMGs were also differentially methylated in their offspring (53 in females, 72 in males) including F2RL3/F2rl3 and RARA/Rara (two of the 6 most significant sites in humans), and the smoking-exposure marker MYOM1/Myom1^66,68^, all of which were replicated in their validation meta-analysis.

A VA-MVP (Veterans Administration - Million Veterans Program) analysis sought to identify causal DNA methylation markers for AAA utilizing an EWAS followed by a Mendelian Randomization (MR) study.^68^ Among the strongest smoking-exposure and AAA-associated CpGs was F2RL3, mentioned above. They also found that hypomethylation near MYOM1 was associated with increased AAA risk. Lastly, of the 23 CpG loci-associated genes they found by MR analysis to be most significantly causative for AAA in humans, 9 were DMGs in both vape-exposed female mice and their male and female offspring in the current study, including Cdk6, Cyp1a1, Ldlr, Nfatc4, Plcl1, Sema7a, Shroom3, Scl24a3 and Wnt6.

Lastly, Hoang et al.^66^ also performed a meta-analysis of human cohorts with DNA methylation measured in blood using Illumina’s EPIC array. They identified 594 CpGs (in 261 genes with murine equivalents) that were differentially methylated with adults exposed to smoking *in utero* (vs. unexposed). Of those, 81 were aortic DMGs in our male offspring and 61 in females.

Notably the gene that historically has displayed the strongest association between smoking and DNA methylation in humans, AHRR, was not found to be a DMG in any of our datasets – possibly identifying a difference between exposure to smoke vs. e-cig vapor.^69^ It is known that the most common human AHRR CpG locations are not conserved in mice^70^, and acute exposure to tobacco smoke, but not e-cig vapor, activates the AHR pathway in male mouse skeletal muscle.^71^

These results suggest significant in-common overlap in DNA methylation processes between humans and mice in response to nicotine exposure, and further that vaping exposure in mothers may lead to durable differential methylation in CpGs of AAA-associated genes in themselves and their offspring, contributing to future disease risk.

In summary, smoking, vaping, and nicotine in particular are potentially capable of not only causing cardiovascular disease, but also altering the epigenome, transmitting disease risk to subsequent generations. This has significant public health implications, as the impact of these substances may persist beyond individual users to their offspring.

There are several limitations to this study. Processing of nicotine is considerably more rapid in mice than in humans, and it is unknown to what extent experimental murine vape exposure actually mimics human use. Further, human genetic diversity likely impacts genomic remodeling responses to toxins, when compared with the uniformity of the genome found in lab mice. In future studies we will examine the impacts of nicotine vape exposure on the uterine environment and germ cells to more clearly determine the mechanisms behind the effects seen in this work, and obtain more detailed data on the specific transcriptional effects of epigenetic changes in offspring. Our experiments provide no data on the effects of paternal exposure, an area which clearly warrants additional study. Lastly, in order to verify that e-cig nicotine exposure leads to sufficient epigenetic germline programming to induce true transgenerational transmission of AAA risk (rather than simply intergenerational), it will be necessary to conduct multi- generational studies, which we intend to pursue going forward.

## ABBREVIATIONS

AAA: abdominal aortic aneurysm
ApoE-/-: Apolipoprotein E knockout mouse strain
AngII: angiotensin II
ATAC-seq: Assay for Transposase-Accessible Chromatin high-throughput sequencing
DAG: differentially accessible gene
DMG: differentially methylated gene
e-cig: electronic cigarette
EWAS: Epigenome-Wide Association Study
F0: parental generation
F1: offspring generation
FRA: F0/F1 female exposed only to room-air
FVF0: F0 female exposed to e-cig vaping
FVF1: F1 female, mother exposed to e-cig vaping
GWAS: Genome-Wide Association Study
MRA: F0/F1 male exposed only to room-air
MVF1: F1 male, mother exposed to e-cig vaping
PPE: porcine pancreatic elastase
RRBS: Reduced Representation Bisulfite high-throughput sequencing
SMC: smooth muscle cell

## Acknowledgments

None

## Supplemental Material

Supplemental Figures 1-10

Supplemental Tables 1-10

