## Supplemental Figures for "Maternal E-cigarette Exposure Augments Murine Abdominal Aortic Aneurysm"

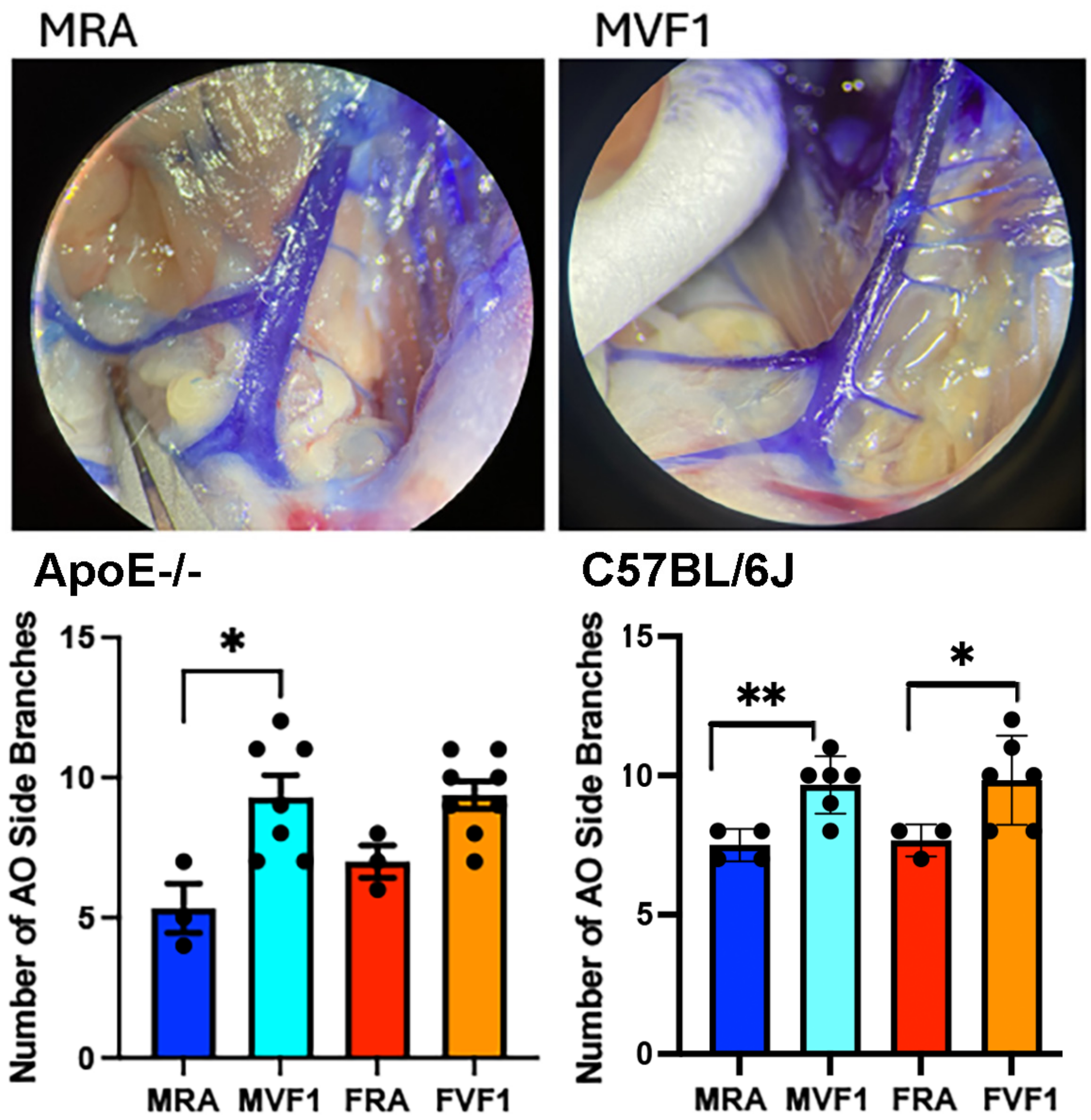


**Supplemental Figure 1:** Aortic side branch proliferation in offspring of e-cig nicotine vaped females (males = MVF1, females = FVF1; Subset B). Top, representative bromophenol-blue-injected abdominal aortae, ApoE-/- males. Bottom (Left ApoE-/- mice, Right C57BL/6J mice), adult offspring display an increased number of small suprarenal aortic side-branches compared with room air controls (males = MRA, females = FRA), *p<0.05.

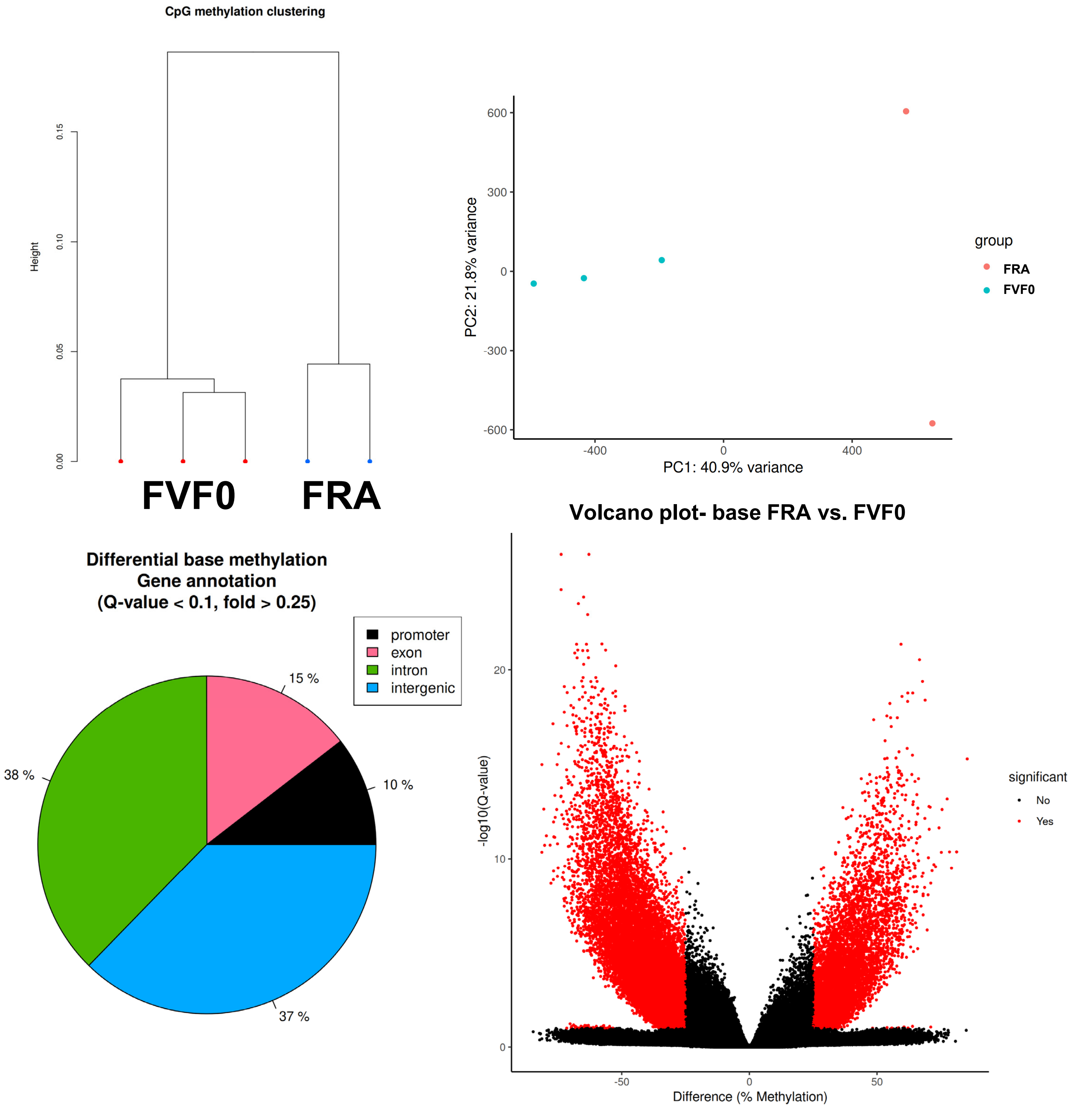


**Supplemental Figure 2: FRA vs. FVF0 (RRBS)**

Findings for Subset A RRBS base methylation comparisons between vaped female F0 C57BL/6J mice and room-air controls. Top left, hierarchical clustering (Pearson). Top right, Principal components analysis (PCA). Bottom left, % locations of differential bases. Bottom right, volcano plot of differential bases.

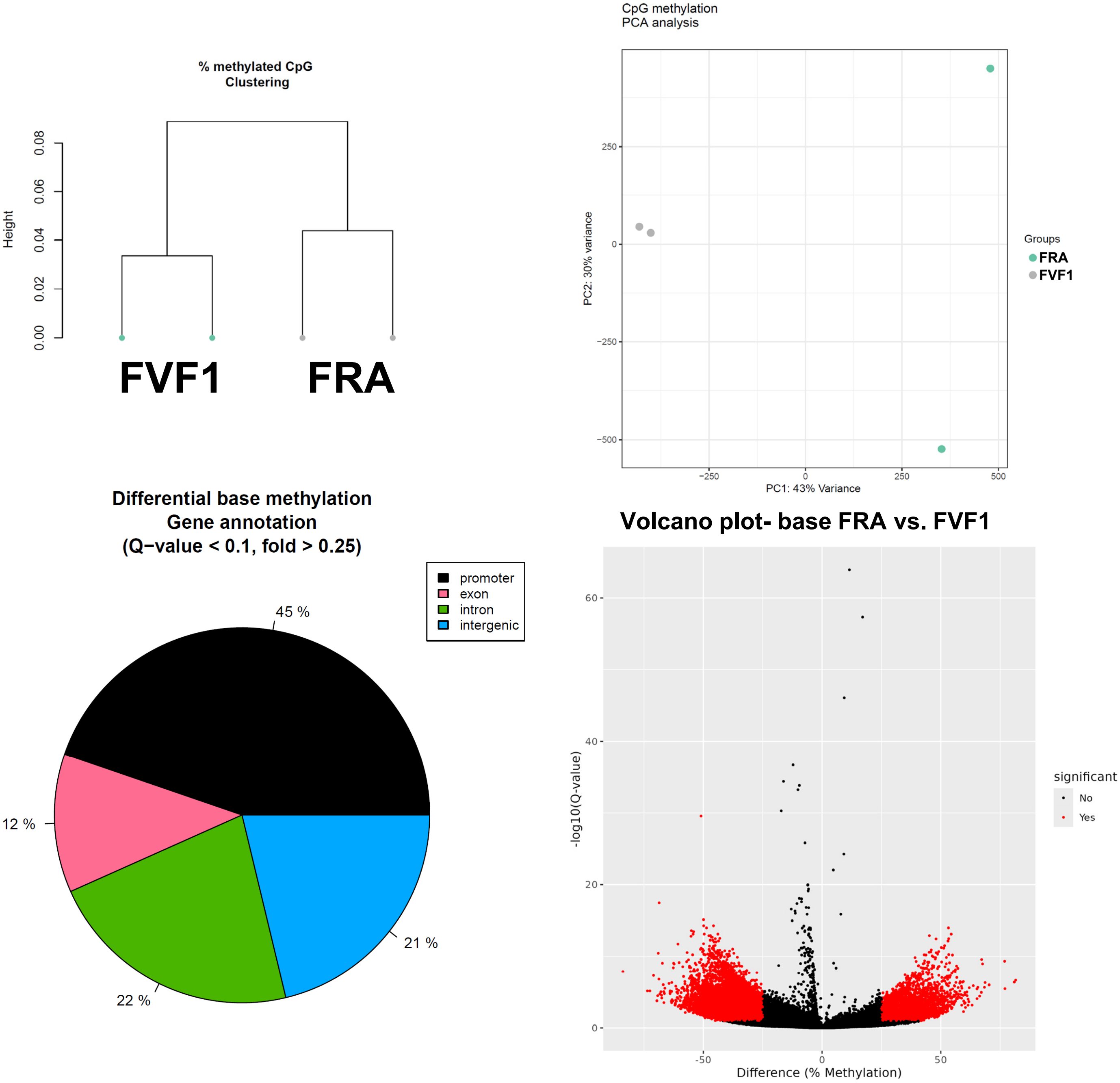


**Supplemental Figure 3: FRA vs. FVF1 (RRBS)**

Findings for Subset B RRBS base methylation comparisons between female C57BL/6J offspring of vaped parent and room-air controls. Top left, hierarchical clustering (Pearson). Top right, Principal components analysis (PCA). Bottom left, % locations of differential bases. Bottom right, volcano plot of differential bases.

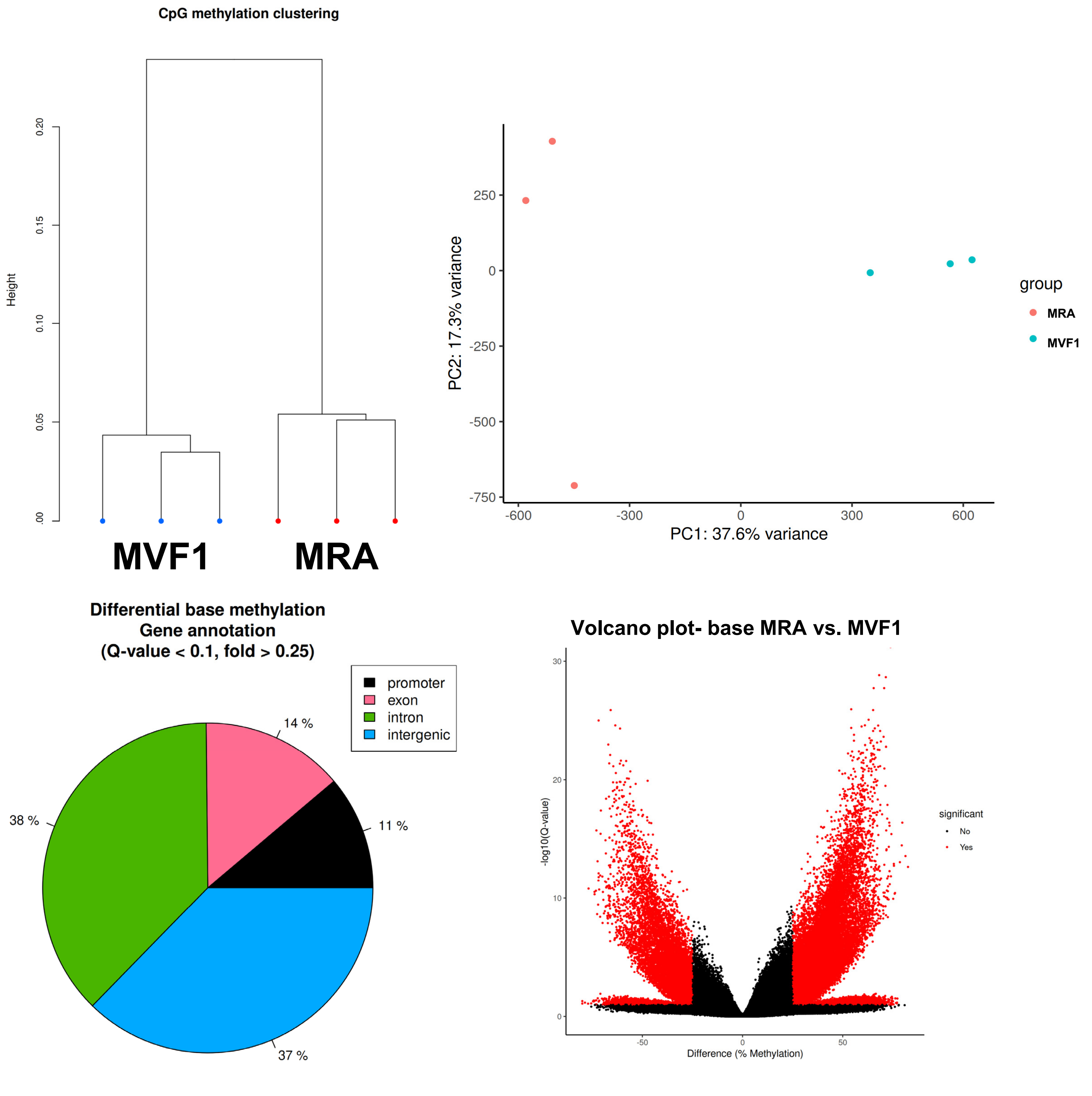


**Supplemental Figure 4: MRA vs. MVF1 (RRBS)**

Findings for Subset B RRBS base methylation comparisons between male C57BL/6J offspring of vaped parent and room-air controls. Top left, hierarchical clustering (Pearson). Top right, Principal components analysis (PCA). Bottom left, % locations of differential bases. Bottom right, volcano plot of differential bases.

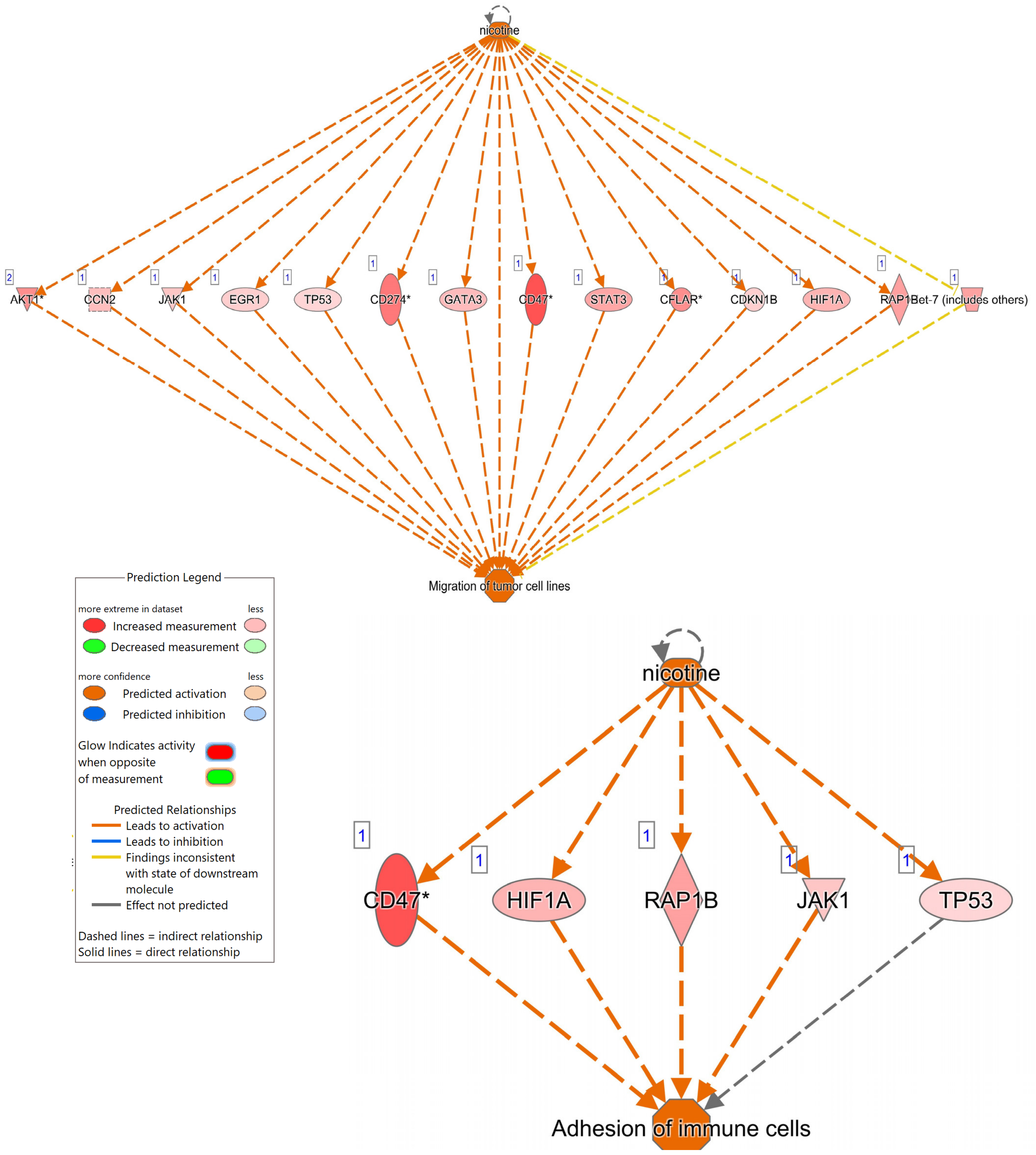


**Supplemental Figure 5a.** IPA Analysis of more accessible (UP) DAGs with e-cig vaping in C57BL/6J females (FRA vs. FVF0 – Subset A). Upstream regulator effect networks were constructed and the networks with the top two consistency ranks are shown (#1 at top with a consistency score of 3.742, and #2 bottom). Consistency score is a measure of how causally consistent and densely connected a regulator effects network is. Prediction Legend shown.

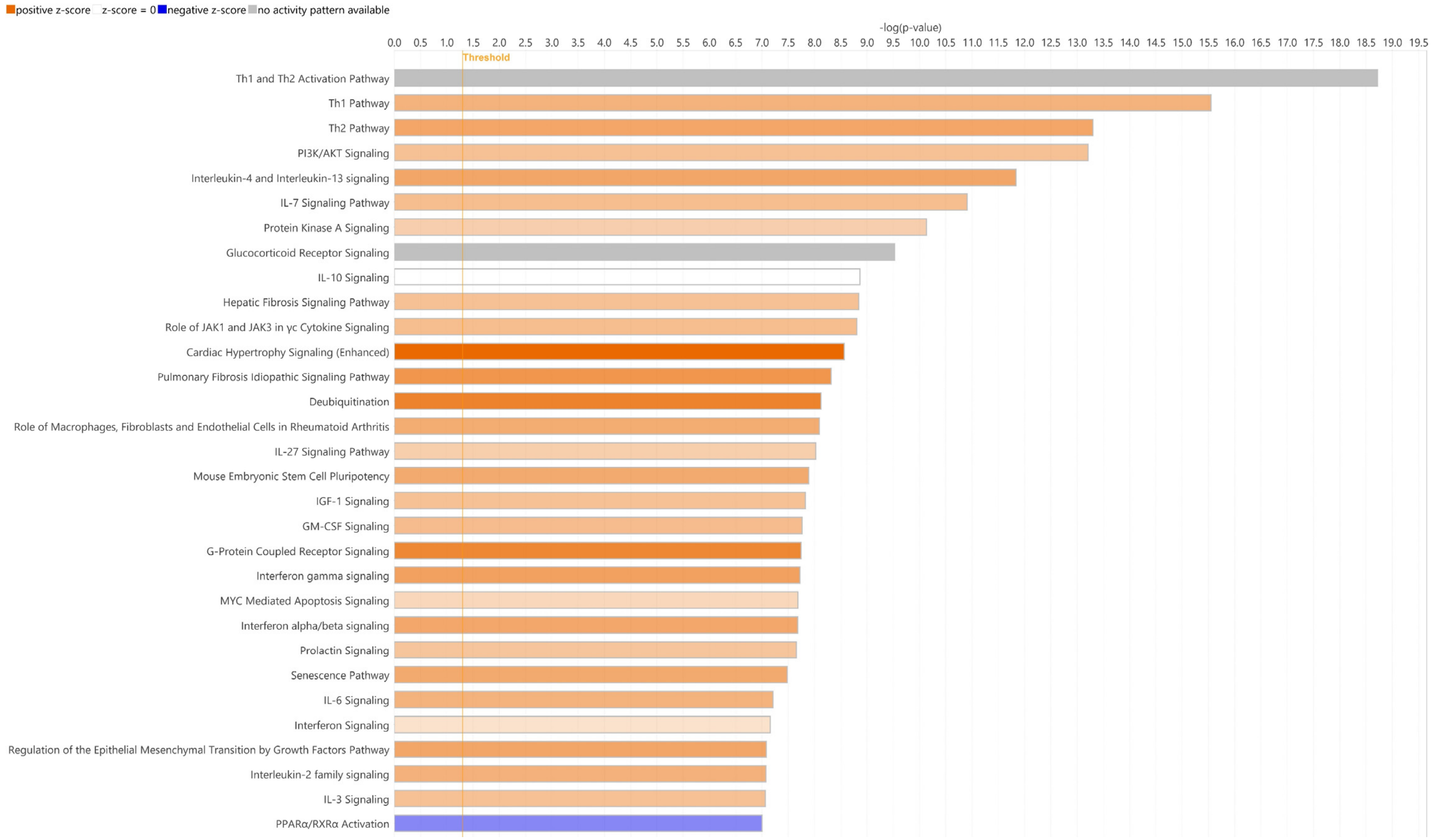


**Supplemental Figure 5b.** IPA Analysis of more accessible (UP) DAGs with e-cig vaping in C57BL/6J females (FRA vs. FVF0 – Subset A). Canonical pathway analysis shown.

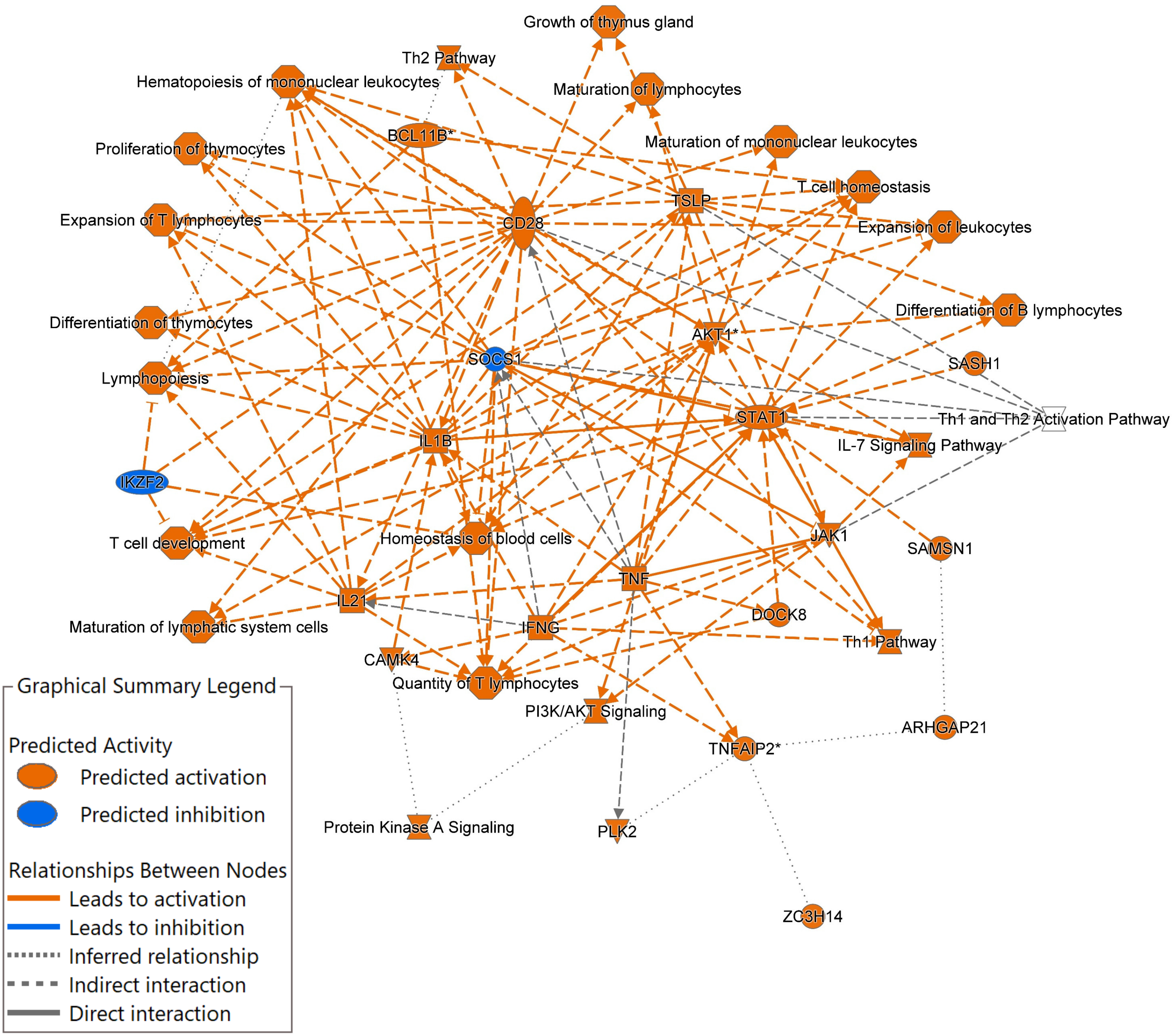


**Supplemental Figure 5c.** IPA Analysis of more accessible (UP) DAGs with e-cig vaping in C57BL/6J females (FRA vs. FVF0 – Subset A). Summary Graph network shown.

#### Supplemental Figure 6a. DAG Motifs – UP peaks FRA vs. FVF0 (Subset A, C57BL/6J)

HOMER de novo Enriched Motifs (note: links are not live)

| Rank | Motif | P-value | log P-pvalue | % of Targets | % of Background | STD(Bg STD) | Best Match/Details |
|---|---|---|---|---|---|---|---|
| 1 | 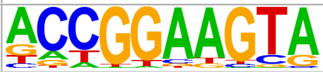 | 1e-21 | -4.997e+01 | 10.21% | 3.46% | 55.4bp (64.9bp) | ELK3/MA0759.1/Jaspar(0.946) More Information \| Similar Motifs Found |
| 2 | 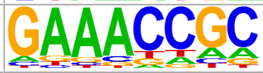 | 1e-13 | -3.195e+01 | 23.51% | 14.54% | 55.3bp (65.0bp) | IRF4(IRF)/GM12878-IRF4-ChIP-Seq(GSE32465)/Homer(0.842) More Information \| Similar Motifs Found |
| 3 | 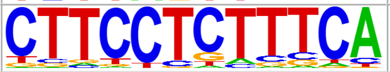 | 1e-12 | -2.934e+01 | 6.36% | 2.25% | 51.6bp (61.9bp) | IKZF1/MA1508.1/Jaspar(0.797) More Information \| Similar Motifs Found |

| Rank | Motif | P-value | log P-pvalue | % of Targets | % of Background | STD(Bg STD) | Best Match/Details |
|---|---|---|---|---|---|---|---|
| 1 | 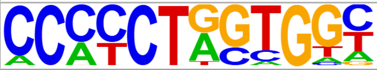 | 1e-99 | -2.294e+02 | 13.97% | 0.77% | 56.4bp (59.7bp) | BORIS(Zf)/K562-CTCFL-ChIP-Seq(GSE32465)/Homer(0.893) More Information \| Similar Motifs Found |
| 2 | 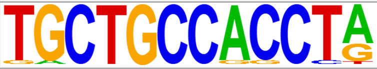 | 1e-23 | -5.489e+01 | 2.37% | 0.06% | 45.0bp (60.1bp) | Unknown3/Drosophila-Promoters/Homer(0.722) More Information \| Similar Motifs Found |
| 3 | 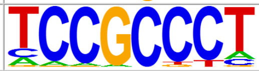 | 1e-17 | -4.116e+01 | 25.69% | 13.95% | 54.4bp (66.9bp) | btd/MA0443.1/Jaspar(0.860) More Information \| Similar Motifs Found |
| 4 | 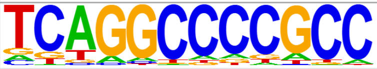 | 1e-14 | -3.340e+01 | 8.98% | 3.11% | 52.8bp (62.7bp) | Sp1(Zf)/Promoter/Homer(0.801) More Information \| Similar Motifs Found |
| 5 | 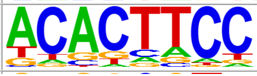 | 1e-13 | -3.193e+01 | 38.90% | 26.50% | 56.4bp (60.9bp) | PU.1(ETS)/ThioMac-PU.1-ChIP-Seq(GSE21512)/Homer(0.851) More Information \| Similar Motifs Found |

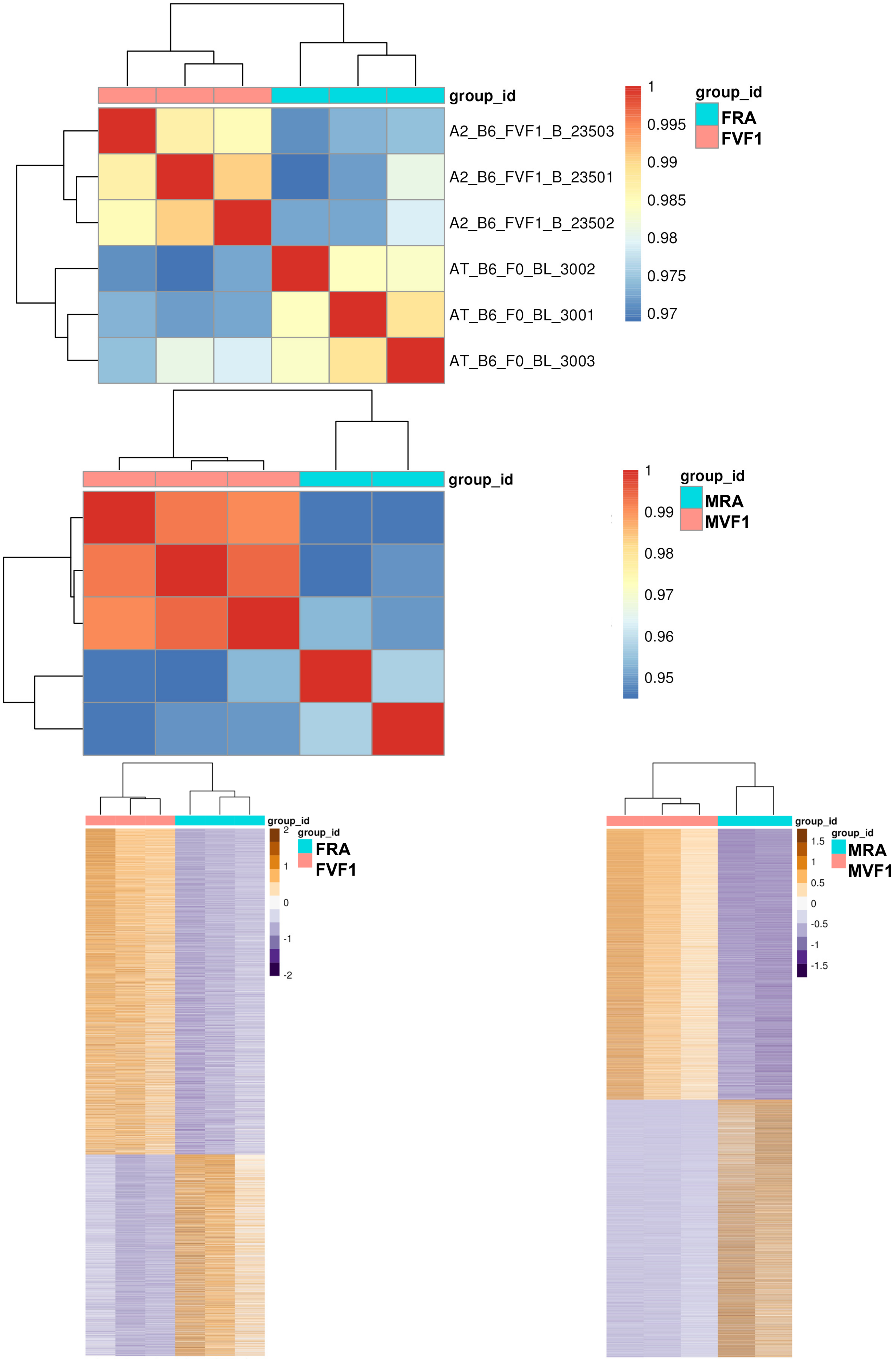


**Supplemental Figure 7.** Impact of maternal vaping on F1-generation adult C57BL/6J female and male mice vs. FRA/MRA controls, blood ATAC-Seq. Analysis identified numerous differential accessibility genes (DAGs, pAdj<0.05, logFC >0.35). Female read range 82-110 million; FRIP score range 16-47%. DAGs FVF1 vs. FRA (2,857 UP and 2,493 DOWN). Male read range 82-95 million; FRIP score range 16-60%. DAGs MVF1 vs. MRA (7,417 UP and 5,977 DOWN). Top, sample PCA Pearson Correlations; Bottom, heatmaps of all significantly regulated DAGs (blue = room air controls, pink = maternal e-cig vaped).

### Supplemental Figure 8a. DAG Motifs – UP peaks FRA vs. FVF1 (Subset B, C57BL/6J)

#### HOMER de novo Enriched Motifs (note: links are not live)

| Rank | Motif | P-value | log P-pvalue | % of Targets | % of Background | STD(Bg STD) | Best Match/Details |
|---|---|---|---|---|---|---|---|
| 1 | 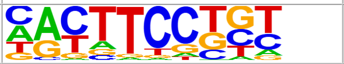 | 1e-200 | -4.622e+02 | 23.57% | 9.96% | 55.2bp (62.9bp) | ETS1(ETS)/Jurkat-ETS1-ChIP-Seq(GSE17954)/Homer(0.947) More Information \| Similar Motifs Found |
| 2 | 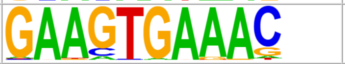 | 1e-66 | -1.522e+02 | 5.05% | 1.55% | 55.1bp (66.5bp) | PU.1-IRF(ETS:IRF)/Bcell-PU.1-ChIP-Seq(GSE21512)/Homer(0.945) More Information \| Similar Motifs Found |
| 3 | 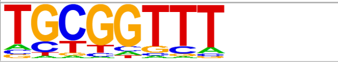 | 1e-51 | -1.194e+02 | 31.75% | 23.07% | 55.2bp (65.1bp) | RUNX2/MA0511.2/Jaspar(0.921) More Information \| Similar Motifs Found |
| 4 | 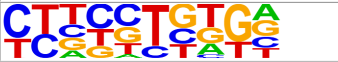 | 1e-36 | -8.315e+01 | 30.03% | 22.89% | 55.8bp (60.8bp) | Ets1-distal(ETS)/CD4+-PolII-ChIP-Seq(Barski_et_al.)/Homer(0.670) More Information \| Similar Motifs Found |
| 5 | 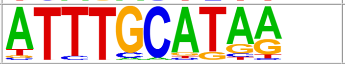 | 1e-32 | -7.456e+01 | 3.21% | 1.18% | 56.3bp (58.4bp) | Oct4(POU,Homeobox)/mES-Oct4-ChIP-Seq(GSE11431)/Homer(0.968) More Information \| Similar Motifs Found |
| 6 | 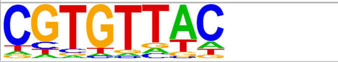 | 1e-29 | -6.899e+01 | 56.25% | 48.79% | 55.2bp (64.6bp) | caup/MA0217.1/Jaspar(0.814) More Information \| Similar Motifs Found |
| 7 | 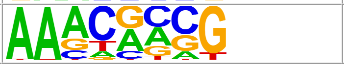 | 1e-29 | -6.695e+01 | 57.58% | 50.24% | 56.8bp (65.8bp) | bin/dmmpmm(Pollard)/fly(0.709) More Information \| Similar Motifs Found |
| 8 | 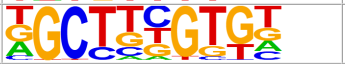 | 1e-24 | -5.665e+01 | 17.54% | 12.82% | 56.6bp (63.0bp) | POL010.1_DCE_S_III/Jaspar(0.709) More Information \| Similar Motifs Found |
| 9 | 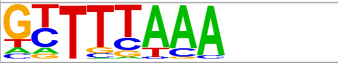 | 1e-23 | -5.482e+01 | 27.50% | 21.87% | 56.0bp (67.0bp) | CHR(?)/Hela-CellCycle-Expression/Homer(0.874) More Information \| Similar Motifs Found |
| 10 | 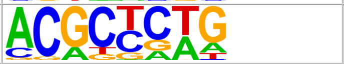 | 1e-17 | -4.010e+01 | 16.81% | 12.90% | 56.2bp (66.5bp) | CG33714(RRM)/Drosophila_melanogaster-RNCMPT00009-PBM/HughesRNA(0.758) More Information \| Similar Motifs Found |
| 11 | 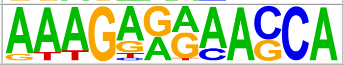 | 1e-16 | -3.887e+01 | 19.87% | 15.71% | 52.3bp (62.7bp) | IRF2/MA0051.1/Jaspar(0.718) More Information \| Similar Motifs Found |
| 12 | 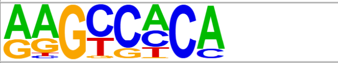 | 1e-16 | -3.847e+01 | 29.01% | 24.21% | 56.7bp (60.5bp) | MET32/MA0334.1/Jaspar(0.801) More Information \| Similar Motifs Found |
| 13 | 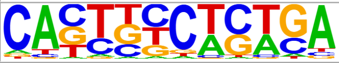 | 1e-14 | -3.307e+01 | 1.02% | 0.31% | 50.3bp (56.1bp) | PU.1(ETS)/ThioMac-PU.1-ChIP-Seq(GSE21512)/Homer(0.724) More Information \| Similar Motifs Found |
| 14 | 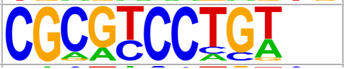 | 1e-12 | -2.921e+01 | 0.41% | 0.06% | 51.3bp (68.3bp) | SWI6(MacIsaac)/Yeast(0.768) More Information \| Similar Motifs Found |

### Supplemental Figure 8b. DAG Motifs – DOWN peaks FRA vs. FVF1 (Subset B, C57BL/6J)

| Rank | Motif | P-value | log P-pvalue | % of Targets | % of Background | STD(Bg STD) | Best Match/Details |
|---|---|---|---|---|---|---|---|
| 1 | 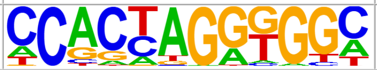 | 1e-248 | -5.731e+02 | 8.59% | 0.56% | 55.3bp (59.8bp) | BORIS(Zf)/K562-CTCFL-ChIP-Seq(GSE32465)/Homer(0.906) More Information \| Similar Motifs Found |
| 2 | 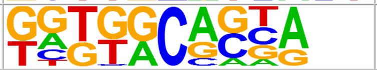 | 1e-29 | -6.692e+01 | 11.22% | 6.21% | 57.8bp (59.0bp) | HIC1(Zf)/Treg-ZBTB29-ChIP-Seq(GSE99889)/Homer(0.693) More Information \| Similar Motifs Found |
| 3 | 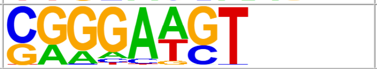 | 1e-18 | -4.154e+01 | 13.18% | 8.77% | 55.0bp (57.5bp) | PB0058.1_Sfpi1_1/Jaspar(0.826) More Information \| Similar Motifs Found |
| 4 | 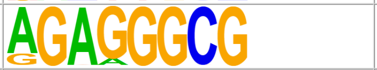 | 1e-17 | -4.010e+01 | 4.45% | 2.09% | 58.3bp (61.9bp) | btd/MA0443.1/Jaspar(0.802) More Information \| Similar Motifs Found |
| 5 | 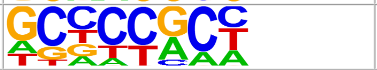 | 1e-13 | -3.190e+01 | 26.41% | 21.10% | 55.7bp (61.6bp) | rst2/MA1431.1/Jaspar(0.796) More Information \| Similar Motifs Found |

### Supplemental Figure 9a. DAG Motifs – UP peaks MRA vs. MVF1 (Subset B, C57BL/6J)

HOMER de novo Enriched Motifs (note: links are not live)

| Rank | Motif | P-value | log P-pvalue | % of Targets | % of Background | STD(Bg STD) | Best Match/Details |
|---|---|---|---|---|---|---|---|
| 1 | 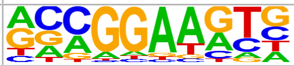 | 1e-422 | -9.721e+02 | 23.60% | 9.55% | 53.9bp (67.2bp) | Fli1(ETS)/CD8-FLI-ChIP-Seq(GSE20898)/Homer(0.974) More Information \| Similar Motifs Found |
| 2 | 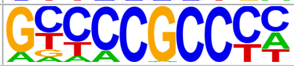 | 1e-172 | -3.967e+02 | 26.38% | 16.07% | 55.4bp (69.9bp) | Sp2(Zf)/HEK293-Sp2.eGFP-ChIP-Seq(Encode)/Homer(0.959) More Information \| Similar Motifs Found |
| 3 | 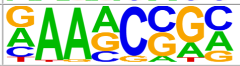 | 1e-121 | -2.794e+02 | 56.18% | 45.23% | 55.7bp (71.4bp) | che-1/MA0260.1/Jaspar(0.834) More Information \| Similar Motifs Found |
| 4 | 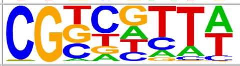 | 1e-112 | -2.587e+02 | 47.58% | 37.23% | 56.1bp (74.0bp) | STB5/MA0392.1/Jaspar(0.793) More Information \| Similar Motifs Found |
| 5 | 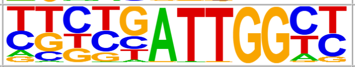 | 1e-93 | -2.151e+02 | 5.27% | 2.03% | 55.1bp (69.6bp) | NFY(CCAAT)/Promoter/Homer(0.959) More Information \| Similar Motifs Found |
| 6 | 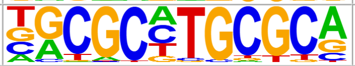 | 1e-84 | -1.945e+02 | 5.26% | 2.14% | 54.0bp (75.2bp) | NRF1/MA0506.1/Jaspar(0.970) More Information \| Similar Motifs Found |
| 7 | 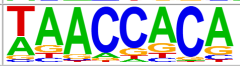 | 1e-81 | -1.869e+02 | 14.81% | 9.23% | 55.3bp (63.5bp) | RUNX(Runt)/HPC7-Runx1-ChIP-Seq(GSE22178)/Homer(0.960) More Information \| Similar Motifs Found |
| 8 | 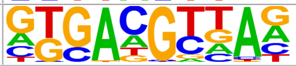 | 1e-80 | -1.852e+02 | 31.05% | 23.27% | 56.0bp (66.1bp) | TGA1(bZIP)/Arabidopsis thaliana/AthaMap(0.843) More Information \| Similar Motifs Found |
| 9 | 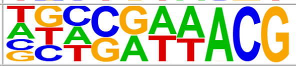 | 1e-67 | -1.543e+02 | 52.06% | 43.98% | 55.2bp (70.8bp) | UPC2/MA0411.1/Jaspar(0.754) More Information \| Similar Motifs Found |
| 10 | 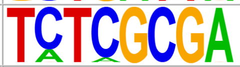 | 1e-59 | -1.378e+02 | 1.40% | 0.27% | 54.0bp (76.5bp) | ZBTB33/MA0527.1/Jaspar(0.898) More Information \| Similar Motifs Found |
| 11 | 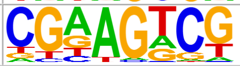 | 1e-43 | -1.000e+02 | 56.12% | 49.64% | 55.9bp (72.3bp) | AT5G22990(C2H2)/col-AT5G22990-DAP-Seq(GSE60143)/Homer(0.726) More Information \| Similar Motifs Found |
| 12 | 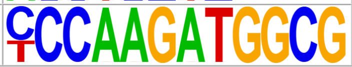 | 1e-17 | -4.083e+01 | 0.21% | 0.02% | 46.6bp (29.5bp) | YY1(Zf)/Promoter/Homer(0.845) More Information \| Similar Motifs Found |

### Supplemental Figure 9b. DAG Motifs – DOWN peaks MRA vs. MVF1 (Subset B, C57BL/6J)

| Rank | Motif | P-value | log P-pvalue | % of Targets | % of Background | STD(Bg STD) | Best Match/Details |
|---|---|---|---|---|---|---|---|
| 1 | 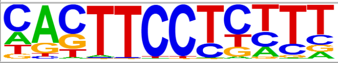 | 1e-340 | -7.838e+02 | 20.77% | 8.51% | 53.1bp (58.5bp) | SpiB(ETS)/OCILY3-SPIB-ChIP-Seq(GSE56857)/Homer(0.951) More Information \| Similar Motifs Found |
| 2 | 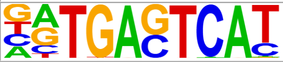 | 1e-240 | -5.534e+02 | 13.44% | 5.12% | 53.0bp (61.5bp) | FOS::JUNB/MA1134.1/Jaspar(0.987) More Information \| Similar Motifs Found |
| 3 | 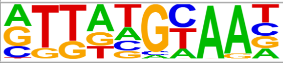 | 1e-182 | -4.198e+02 | 14.99% | 6.98% | 54.6bp (60.7bp) | MF0006.1_bZIP_cEBP-like_subclass/Jaspar(0.922) More Information \| Similar Motifs Found |
| 4 | 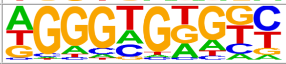 | 1e-130 | -3.013e+02 | 19.23% | 11.22% | 55.7bp (58.3bp) | KLF5(Zf)/LoVo-KLF5-ChIP-Seq(GSE49402)/Homer(0.936) More Information \| Similar Motifs Found |
| 5 | 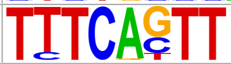 | 1e-85 | -1.973e+02 | 10.66% | 5.78% | 55.5bp (62.0bp) | IRF8(IRF)/BMDM-IRF8-ChIP-Seq(GSE77884)/Homer(0.868) More Information \| Similar Motifs Found |
| 6 | 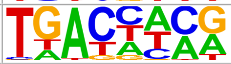 | 1e-61 | -1.416e+02 | 37.64% | 30.20% | 55.5bp (61.0bp) | WRKY48/MA1088.1/Jaspar(0.772) More Information \| Similar Motifs Found |
| 7 | 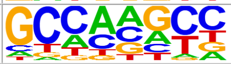 | 1e-53 | -1.235e+02 | 36.84% | 29.92% | 56.1bp (61.5bp) | RIM101/MA0368.1/Jaspar(0.798) More Information \| Similar Motifs Found |
| 8 | 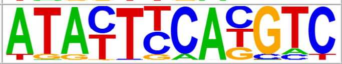 | 1e-37 | -8.545e+01 | 0.22% | 0.01% | 57.1bp (51.8bp) | ABF4/MA1659.1/Jaspar(0.623) More Information \| Similar Motifs Found |
| 9 | 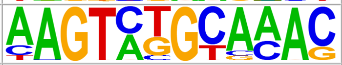 | 1e-22 | -5.077e+01 | 0.17% | 0.01% | 55.7bp (35.9bp) | Oct11(POU,Homeobox)/NCIH1048-POU2F3-ChIP-seq(GSE115123)/Homer(0.638) More Information \| Similar Motifs Found |
| 10 | 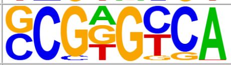 | 1e-20 | -4.704e+01 | 29.22% | 25.26% | 56.0bp (62.7bp) | PB0153.1_Nr2f2_2/Jaspar(0.766) More Information \| Similar Motifs Found |

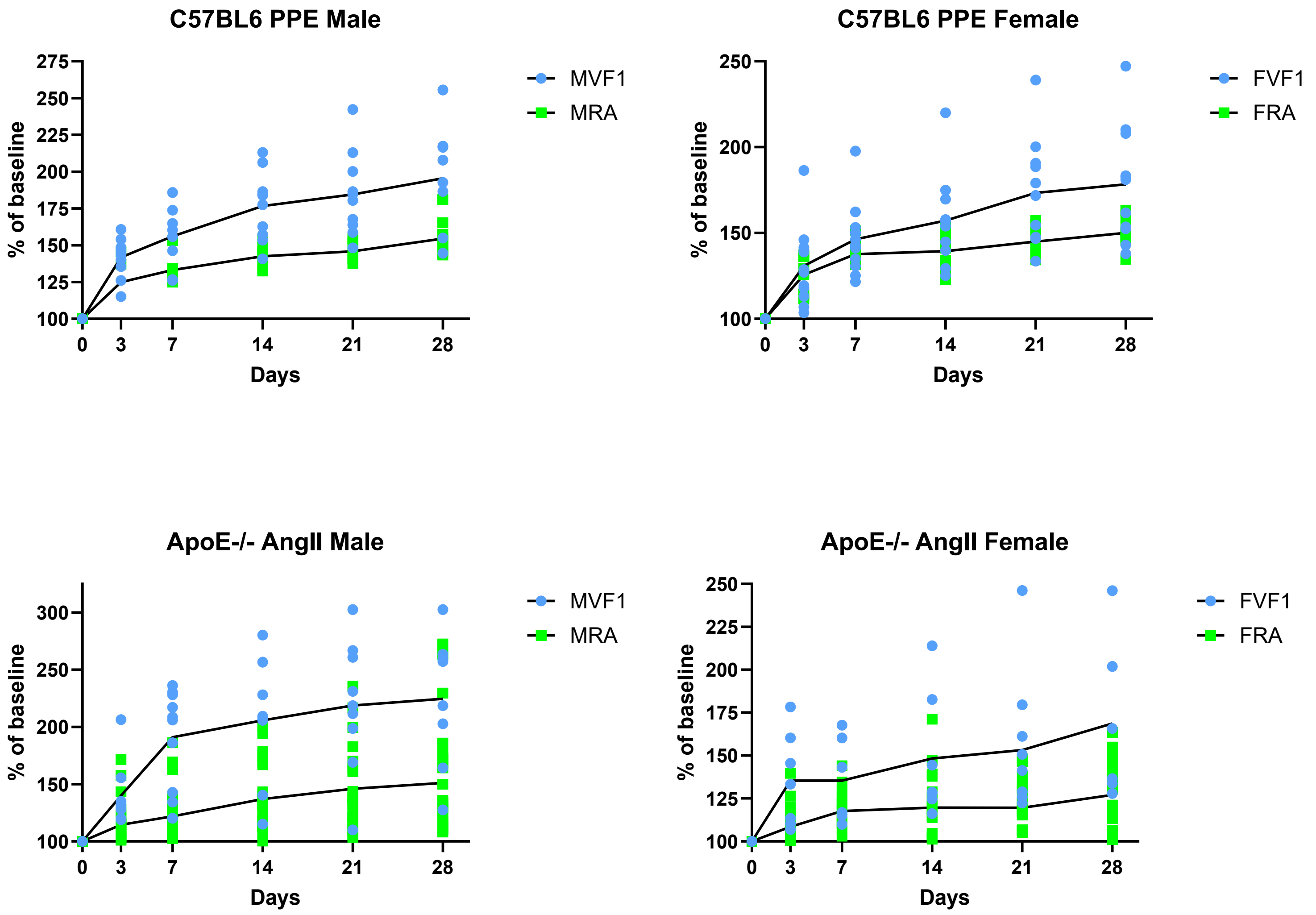


**Supplemental Figure 10**. Alternative format of Figures 3a-d, all data points shown.
